# Transcriptional Regulation of Protein Synthesis by Mediator Kinase in MYC-driven Medulloblastoma

**DOI:** 10.1101/2024.03.08.584103

**Authors:** Dong Wang, Caitlin Ritz, Angela Pierce, Breauna Brunt, Yuhuan Luo, Nathan Dahl, Sujatha Venkataraman, Etienne Danis, Kamil Kuś, Milena Mazan, Tomasz Rzymski, Bethany Veo, Rajeev Vibhakar

## Abstract

MYC-driven medulloblastoma (MB) is a highly aggressive cancer type with poor prognosis and limited treatment options. Through CRISPR-Cas9 screening across MB cell lines, we identified the Mediator-associated kinase CDK8 as the top dependence for MYC-driven MB. Loss of CDK8 markedly reduces MYC expression and impedes MB growth. Mechanistically, we demonstrate that CDK8 depletion suppresses ribosome biogenesis and mRNA translation. CDK8 regulates occupancy of phospho-Polymerase II at specific chromatin loci facilitating an epigenetic alteration that promotes transcriptional regulation of ribosome biogenesis. Additionally, CDK8-mediated phosphorylation of 4EBP1 plays a crucial role in initiating eIF4E-dependent translation. Targeting CDK8 effectively suppresses cancer stem and progenitor cells, characterized by increased ribosome biogenesis activity. We also report the synergistic inhibition of CDK8 and mTOR *in vivo* and *in vitro*. Overall, our findings establish a connection between transcription and translation regulation, suggesting a promising therapeutic approach targets multiple points in the protein synthesis network for MYC-driven MB.

## Introduction

Medulloblastoma (MB) is the most common malignant pediatric tumor, accounting for 15-20% of childhood brain tumors^1^. Molecular profiling and genetic analysis initially categorized medulloblastoma (MB) into four subgroups: WNT, SHH, Group 3, and Group 4^2,3^. Among these groups, patients with MYC-driven Group 3 MB (G3-MB) commonly experience relapse accompanied by metastatic spread and local recurrence, resulting in long-term survival rates of less than 5%^4^. To date, targeted options for G3-MB tumors are lacking, in part because of the incomplete understanding of tumorigenic mechanisms.

Dysregulated expression of the MYC proto-oncogene contributes to the development of most human tumors^5^. Numerous studies have demonstrated that MYC plays a pivotal role in the regulation of protein synthesis^6–9^. MYC affects cell proliferation, growth, and nucleolar size, and is associated with marked changes in the total rate of protein synthesis^10^. It also regulates ribosome biogenesis either directly by upregulating ribosomal RNA and protein components through chromatin structure remodeling, or indirectly by controlling essential auxiliary factors involved in rRNA processing, ribosome assembly, and subunit transportation from the nucleus to the cytoplasm^6,8,11–13^. Chromatin remodeling is an essential aspect of these processes through which MYC directly activates RNA polymerases^14–16^. Understanding the dysregulated protein synthesis in MYC-driven oncogenesis is crucial for developing targeted therapeutic interventions that leverage the inherent vulnerabilities of these pathways in the context of tumor development.

The Mediator Kinase cyclin-dependent kinase 8 (CDK8) has been demonstrated to maintain stemness and tumorigenicity by regulating the MYC pathway^17^. CDK8 associates with the mediator complex, which is a large multi-subunit complex that regulates transcription by connecting enhancer-bound transcription factors to RNA polymerase II^18,19^. Elevated expression of CDK8 has been linked to shorter relapse-free survival in various types of cancer, including colon cancer, breast cancer, GBM, and hepatocellular carcinoma^17,20–22^. In this study, we found that CDK8 was one of the most essential genes for MB. Importantly, CDK8 depletion suppressed protein synthesis, suggesting its cooperation with MYC to drive tumorigenesis. We demonstrated that CDK8 depletion substantially reduced MYC expression and induced pronounced transcriptional changes, consequently resulting in the suppression of ribosomal gene expression, and impeding the growth of MYC-driven MB. Furthermore, CDK8 inhibition with a novel inhibitor, RVU120, synergized with mTOR inhibition to suppress MYC-driven MB. This work holds the promise of significantly advancing the understanding of MYC-driven oncogenesis and provides critical preclinical data essential for the development of novel therapies targeting CDK8 and mTOR in MYC-driven medulloblastoma.

## Results

### Targeting CDK8 suppresses medulloblastoma progression

To systematically identify genes representing therapeutic vulnerabilities in MYC-driven MB, we previously performed CRISPR-Cas9 screening with a specific targeting of 1140 druggable genes across three MYC-amplified human G3-MB cell lines^23,24^. Further analysis identified CDK8 as an essential gene for MB tumor growth (Fig. 1a and Extended Data Fig. 1a,b). We next explored the significance of CDK8 by leveraging the Cancer Dependency Map (DepMap), a platform that utilizes the CRISPR gene knockout to map gene dependencies across hundreds of cancer types^25^. CDK8 is essential for various types of cancers, with medulloblastoma being the most enriched cancer type (Fig. 1b). Within medulloblastoma, CDK8 stands out as a top dependence, similar to OTX2, Neurog1, and Neurod1, well-established genes that sustain stemness and drive proliferation exclusively in medulloblastoma (Fig. 1c)^26–29^. We found that CRISPR-Cas9-mediated CDK8 knockout suppressed MB cell proliferation (Fig. 1d). Furthermore, CDK8 is the only gene with clinically relevant inhibitors among the top dependencies (CDK8, RALA, Neurog1, Neurod1, and OTX2), making it a potential target for the treatment of G3-MB.

**Fig. 1.**
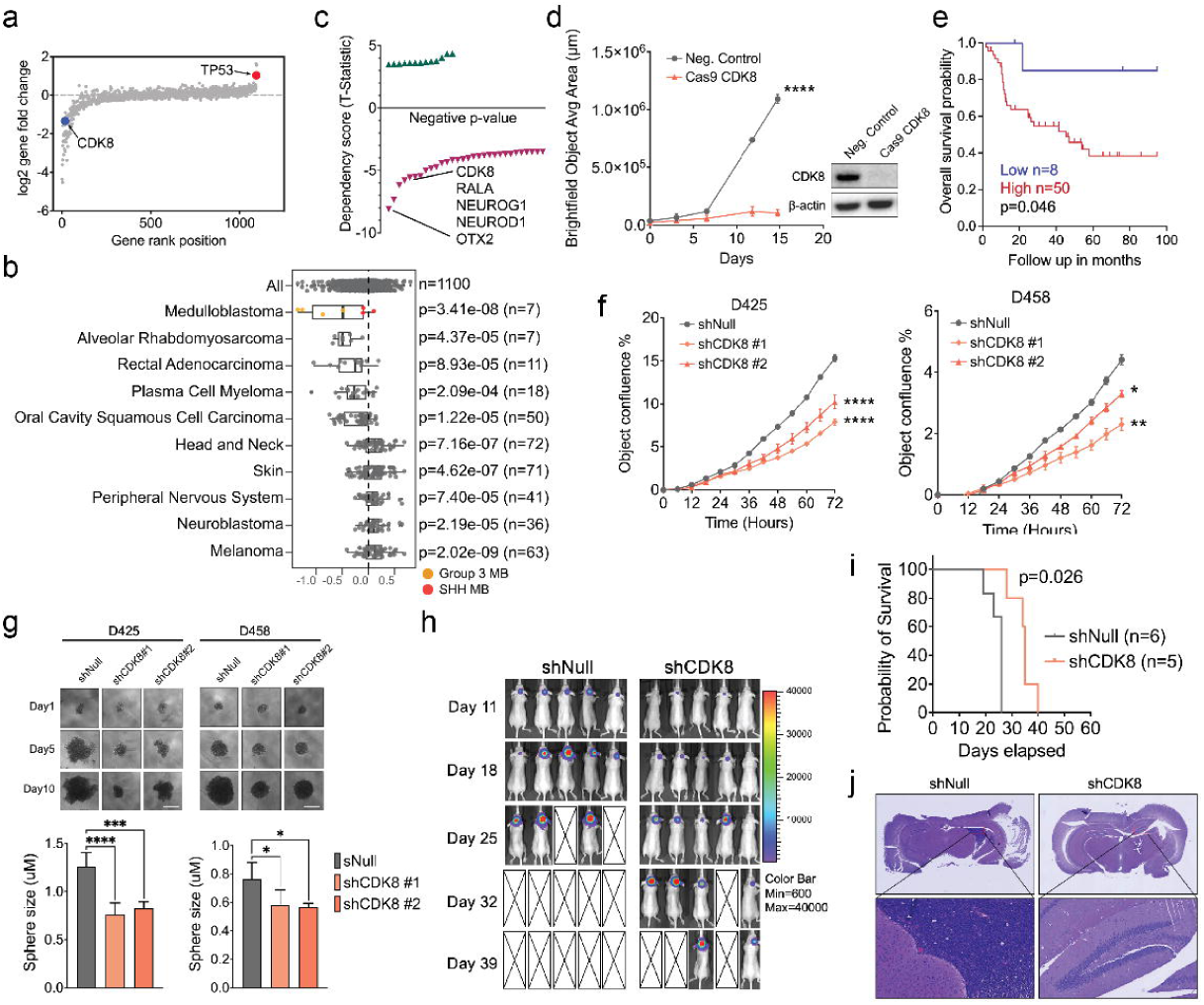
| Targeting CDK8 suppresses medulloblastoma progression. **a**, Log-fold change of 1140 druggable gene expression in CRISPR-Cas9 screening across D341, D425, and D458 cell lines. CDK8 is a negative selective gene that is essential for MB tumor growth. TP53 is labeled with red as positive control. **b,** The enriched lineages plot from DepMap indicates the DEMETER2 score of CDK8 across various cancer types. A lower DEMETER2 score indicates a higher likelihood that the gene is essential in a given cell line. A score of 0 indicates a gene is not essential; correspondingly -1 is comparable to the median of all pan-essential genes. n= indicates the number of cell lines plotted in that lineage. **c,** The Dependency score from Depmap specific for medulloblastoma. Labels below zero denote genes that are essential for the growth of MB. **d,** CRISPR-Cas9 targeting CDK8 suppressed the growth of D458 cells in non-FBS neurosphere media. n = 5. Mean ± SD. Statistical analysis: One-way ANOVA. **e,** Kaplan-Meier survival curve demonstrating the association between the level of CDK8 and overall survival within the subset of MYC-high expression Group 3 MB patient cases. Statistical analysis: log-rank (Mantel-Cox) test. **f,** Growth of shNull and shCDK8-transduced D425 or D458 cells assayed in Incucyte live-cell analysis system. n = 5. Mean ± SD. Statistical analysis: one-way ANOVA. **g,** Representative images of neurosphere size in CDK8 depleted D425 or D458 MB cell lines at 1, 5, and 10 days are shown. n = 5. Mean ± SD. Scale bar, 400 μm. Statistical analysis: one-way ANOVA. **h,** Representative bioluminescence images of shCDK8 or shNull D458 xenografts. Shown are days post injection. Color scales indicate bioluminescence radiance in photons/sec/cm2/steradian. **i,** Kaplan-Meier survival curve of shCDK8 or shNull D458 xenograft mice. Statistical analysis: log-rank (Mantel-Cox) test. **j,** Representative histological images showing a significant reduction in tumor formation in D458 xenografts generated from the shCDK8 cells compared with the shNull cells. Three mice from each group were sacrificed 20 days after tumor implantation. Original magnification, ×40.

Using single-cell murine cerebellar transcript data, we found relatively low expression of CDK8, and genes associated with the Mediator Complex, suggesting that low levels of mediators may be sufficient for normal developmental processes (Extended Data Fig. 1c). CDK8 exhibited notably higher level in multiple G3-MB cells compared to that in normal cerebellar tissue (Extended Data Fig. 1d). Proteomic analysis of 45 MB patient samples also demonstrated elevated CDK8 protein in G3-MB^30^ (Extended Data Fig. 1e). To further examine CDK8 in MB, we performed an analysis using a cohort of 763 described MB samples, along with normal cerebellar samples. G3-MB expressed higher expression of CDK8 than the normal cerebellum, particularly in subtypes Group 3β and 3γ with elevated c-MYC (high-MYC MB) (Extended Data Fig. 1f, g). Kaplan–Meier survival analysis performed on the same dataset revealed a correlation between CDK8 expression and poor overall survival in high-MYC MB (Fig. 1e).

To determine the dependence of MB on CDK8, we inhibited CDK8 expression in MB cells using lentivirus-mediated CDK8 shRNAs. Loss of CDK8 led to a notable decrease in both cell proliferation and the level of MYC (Fig. 1f and Extended Data Fig. 1h). Additionally, CDK8 knockdown significantly decreased neurosphere growth in MB cells (Fig. 1g). To further examine the *in vivo* effects of CDK8 on tumor formation, D458 MB cells were transduced with either a control shRNA sequence (shNull) or shRNAs targeting CDK8 (shCDK8). Subsequently, these cells were implanted intracranially into immunodeficient mice. Knockdown of CDK8 inhibited tumor growth and prolonged the survival of intracranial tumor-bearing mice relative to shNull, reinforcing CDK8 as a crucial factor governing the growth of MYC-amplified MB (Fig. 1h-j and Extended Data Fig. 1i).

### RVU120 inhibits CDK8 activity in medulloblastoma cells

Several small-molecule inhibitors targeting CDK8 are currently undergoing preclinical development^31–34^. Our evaluation of eight CDK8 selective inhibitors demonstrated a broad range of half-maximal inhibitory concentrations (IC50) across three G3-MB cell lines (Extended Data Fig. 2a). Among them, RVU120 exhibited remarkable potency, with the lowest IC50 values (Fig. 2a). We assessed the IC50 of RVU120 in various MB cell lines and non-transformed normal human astrocytes (NHA). In G3-MB cells, the 72-hour IC50 concentration ranged from 125.90 1509.00 nM. NHA displayed significantly higher resistance to RVU120, with an IC50 concentration of 4349.00 nM (Fig. 2b). Importantly, RVU120 treatment reduced the viability of patient-derived primary G3-MB cells, further confirming the efficacy of RVU120 in treating G3-MB (Fig. 2c and Extended Data Fig. 2b).

**Fig. 2.**
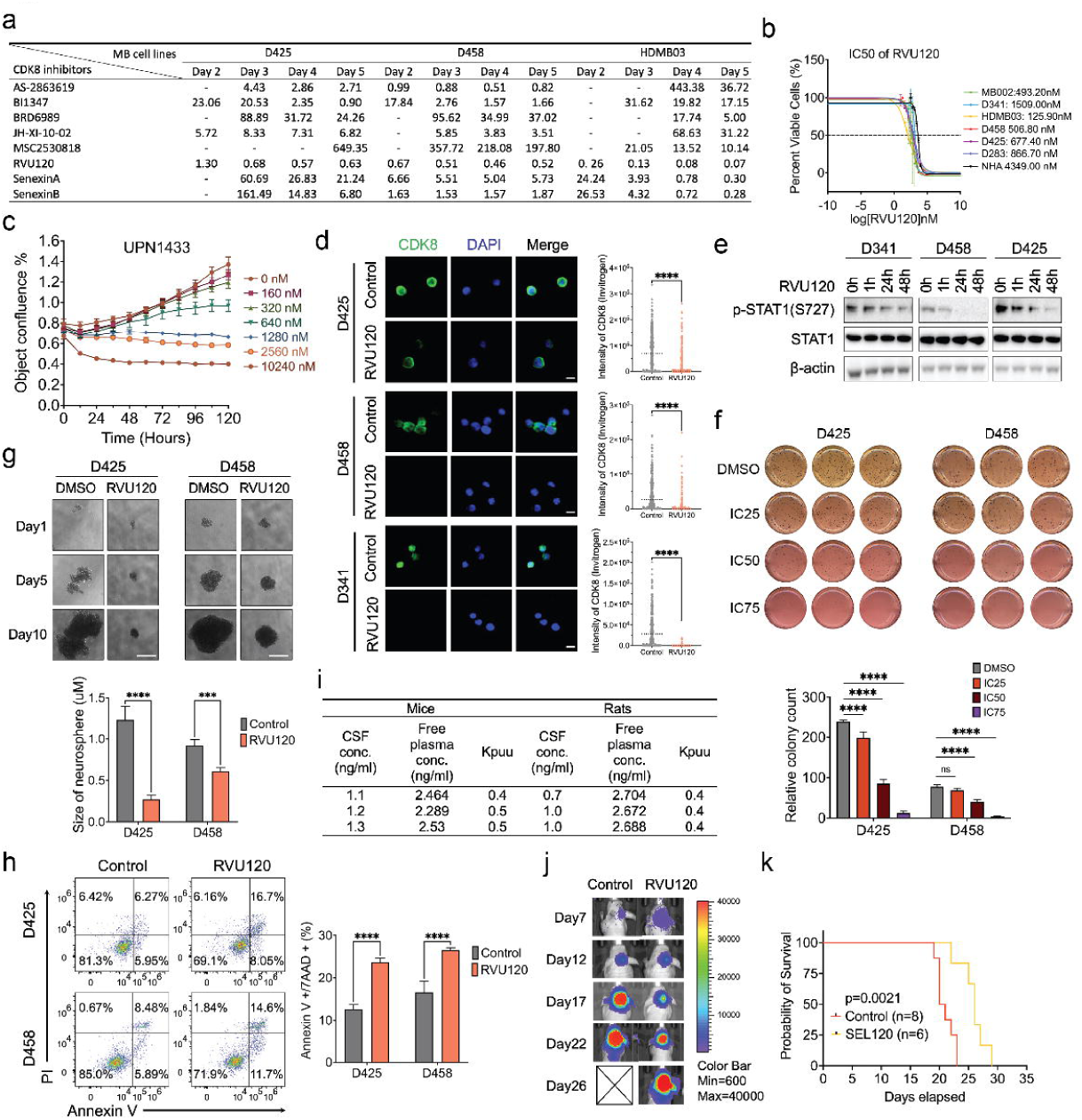
| RVU120 inhibits CDK8 activity in medulloblastoma cells. **a**, Half-maximal inhibitory concentration (IC50) determination of various CDK8 inhibitors in D425, D458, and HDMB03 MB cell lines. Unit: μmol. **b,** IC50 of RVU120 at 72 h in various MB cell lines and normal human astrocytes (NHA) cells. **c,** Dose-dependent proliferation curve of RVU120 treated primary cultured medulloblastoma cells from a G3-MB patient. **d,** Immunofluorescence of CDK8 (green) and DAPI (blue) at 40X magnification. D425, D458, and D341 cells were treated with IC50 RVU120 for 48 h. Scale bar, 10 μm. Statistical analysis: Mann-Whitney Wilcoxon test. **e,** Immunoblot demonstrates time-course analysis of p-STAT1(S727) protein levels with treatment of IC50 RVU120 across D341, D458, D425 MB cell lines. **f,** Methylcellulose assay in D425 and D458 cells treated with DMSO, IC25, IC50, and IC75 for 2 weeks. Statistical analysis: two-way ANOVA. **g,** Representative images of neurosphere size in IC40 RVU120 treated D425 or D458 MB cell lines at 1, 5, and 10 days are shown. n = 5. Mean ± SD. Scale bar, 400 μm. Statistical analysis: one-way ANOVA. **h,** Annexin V apoptosis assay. D425 and D458 cells were treated with IC50 RVU120 for 48 h, stained with Annexin V, and measured by flow cytometry. n = 3. Statistical analysis: student t-test. **i,** The concentration of the compound in the CSF corresponds to the free concentration in the brain. A ratio of less than one in rodents indicates the possibility of intercellular trapping. Kpuu: Unbound partition coefficient. **j,** Representative bioluminescence images of mice treated with RVU120 (40mg/kg, daily, oral gavage) compared with those of the control cohort. **k,** Kaplan–Meier survival curves for animals treated with Control or RVU120. Statistical analysis: log-rank (Mantel-Cox) test.

We observed that treatment with RVU120 led to decreased CDK8 expression and a concurrent reduction in p-STAT1 levels, which is a direct target of CDK8^35^ (Fig. 2d,e and Extended Data Fig. 3a,b). Using a methylcellulose colony-forming assay, we found that CDK8 inhibition suppressed colony formation and neurosphere growth in G3-MB cells (Fig. 2f,g and Extended Data Fig. 3c). Additionally, flow cytometry analysis revealed a substantial increase in the total percentage of apoptotic cells following RVU120 treatment, as determined by both annexin V and active caspase 3 staining using flow cytometry (Fig. 2h and Extended Data Fig. 3d). To assess the potential intracranial efficacy of RVU120 *in vivo*, we evaluated its unbound partition coefficient, which determines the concentration of the compound in the CSF, corresponding to its free concentration in the brain (Fig. 2i). A ratio value of approximately 0.4 was observed, indicating permeation into the brain^36^. Furthermore, in our orthotopic MB xenograft model, we found that administration of RVU120 extended the survival of mice in the treatment group (Fig. 2j,k and Extended Data Fig. 3e). Collectively, these findings reveal an oncogenic role of CDK8 in MB and highlight the therapeutic potential of RVU120 for treatment G3-MB.

### CDK8 depletion leads to repression of mRNA translation and ribosome biogenesis

To understand the mechanisms underlying CDK8 regulation, we performed RNA-Seq of MB cells with genetic knockdown or pharmacological inhibition of CDK8. Enrichment analysis demonstrated that CDK8 depletion altered the hallmark features of MB, including neuronal differentiation, photoreceptor cell maintenance, and nervous system development (Extended Data Fig. 4a). Gene Set Enrichment Analysis (GSEA) revealed a significant alteration in gene sets in CDK8 knockdown cells compare to control cells (Fig. 3a). Notably, many gene sets of Gene Ontology (GO) terms related to mRNA translation were significantly decreased (Fig. 3b). The same effect of CDK8 on mRNA translation was aligned with the pathway alterations observed in RVU120 treated MB cells compared with control cells, further confirming the specific inhibition of CDK8 by RVU120 (Fig. 3c, Extended Data Fig. 4b). To examine the functional role of CDK8 in protein synthesis, we performed an O-propargyl-puromycin (OPP) assay, which involves the introduction of a modified puromycin analog into cells, using click chemistry to visualize and quantify the rates of protein synthesis. Treatment with RVU120 led to a decrease in the OPP signal, from 1h to 48h post-treatment, demonstrating the role of CDK8 in regulating protein synthesis (Fig. 3d).

**Fig. 3.**
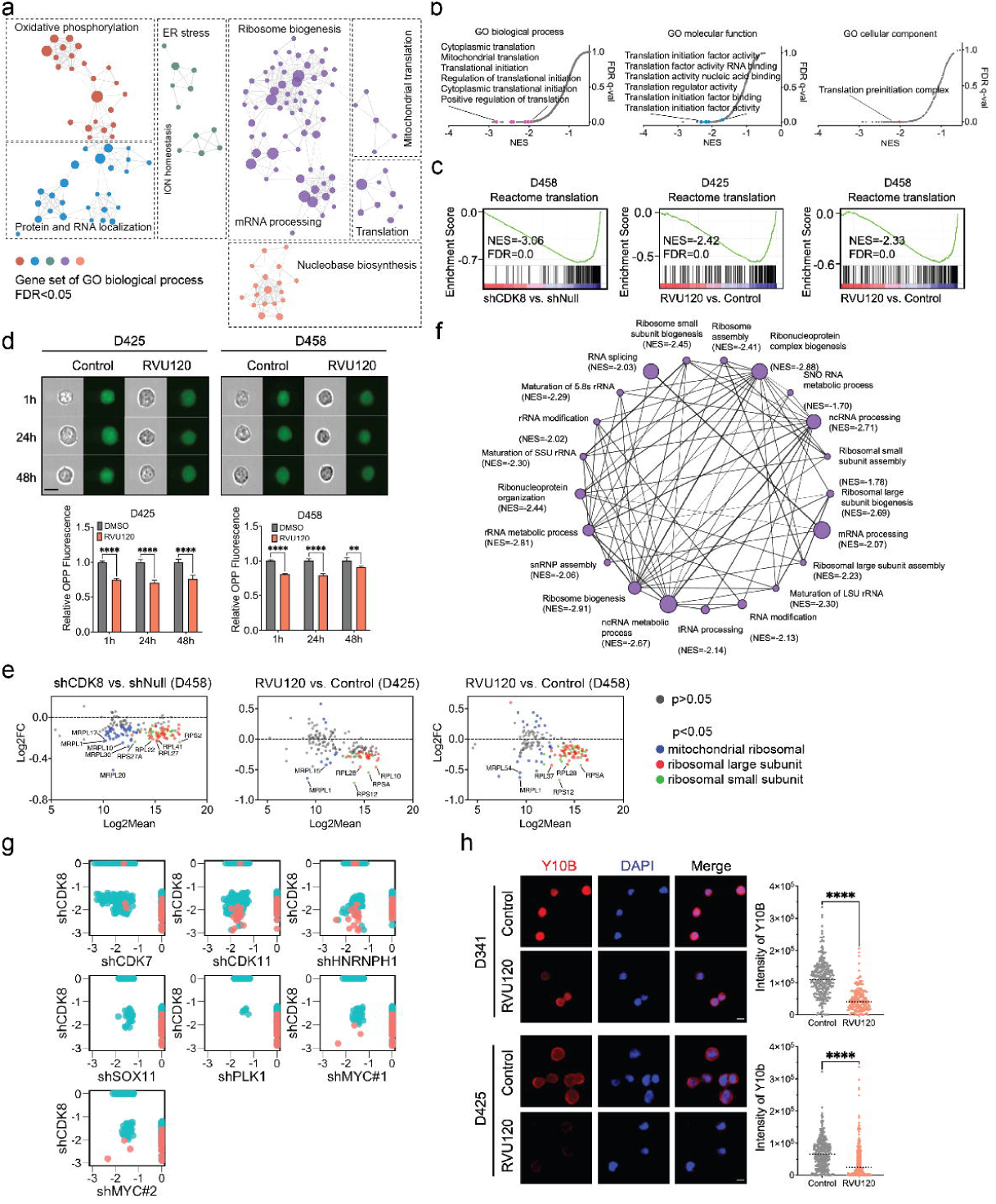
| CDK8 depletion leads to repression of mRNA translation and ribosome biogenesis. **a**, Gene Set Enrichment Analysis (GSEA) indicated alterations in gene sets in D458 cells transfected with shCDK8 compared to cells transfected with shNull. **b,** GSEA revealed significant downregulation of three categories of gene ontology terms related to mRNA translation in shCDK8 D458 cells compared to shNull cells. **c,** GSEA demonstrated downregulation of Reactome pathways related to mRNA translation following genetic knockdown or pharmacological inhibition of CDK8 in MB cells. **d,** O-propargyl-puromycin assay in IC50 RVU120 treated D425 and D458 cells compared with control cells. Scale bar, 20 μm. Statistical analysis: two-way ANOVA. **e,** RNA-Seq demonstrated alterations in the expression of mitochondrial and cytoplasmic ribosomal genes. **f,** The GSEA network revealed downregulation of gene sets associated with ribosome biogenesis in shCDK8 D458 cells compared to shNull D458 cells. The plot was generated using Cytoscape, where each node represents the gene counts within specific gene sets. **g,** GSEA analysis shows GO biological process gene sets (p<0.05) in knockdown of CDK7, CDK11, HNRNPH1, SOX11, and PLK1. CDK8 depletion resulted in a more significant reduction in gene sets related to ribosome biogenesis and mRNA translation compared to other genes. **h,** Immunofluorescence of Y10B (red, anti-ribosomal RNA) and DAPI (blue) at 40X. D425 and D341 cells were treated with the IC50 of RVU120 for 48 h. Scale bar, 10 μm. Statistical analysis: Mann-Whitney Wilcoxon test.

Ribosome biogenesis, which involves the coordinated assembly of ribosomal RNA (rRNA) and ribosomal proteins (RPs), plays a crucial role in regulating mRNA translation by producing functional ribosomes^10^. In MYC-driven cancer cells, including G3-MB cells, ribosomal genes typically demonstrated higher expression levels than other genes (Extended Data Fig. 4c). Upon CDK8 depletion, multiple cytoplasmic and mitochondrial ribosomal genes were downregulated, leading to the significant repression of gene sets associated with ribosomal biogenesis, such as ribosome assembly, ribonucleoprotein complex biogenesis, rRNA maturation, and rRNA modification (Fig. 3e,f and Extended Data Fig. 4d). MYC plays a pivotal role in regulating mRNA translation and is a primary driver of ribosome biogenesis^10^. Therefore, we examined gene set alterations following knockdown of MYC or other related genes (PLK1, CDK7, CDK9, SOX11, and HNRNPH1), all of which are known to affect MYC expression^23,24,37,38^. Remarkably, CDK8 depletion led to a significant decrease in the number of gene sets associated with mRNA translation and ribosome biogenesis, indicating the role of CDK8 in collaborating with MYC to regulate protein synthesis in MB (Fig. 3g). Furthermore, upon RVU120 treatment, we observed a reduction in 5.8S rRNA levels, as indicated by Y10b immunostaining, providing additional evidence to support the impact of CDK8 on ribosome biogenesis (Fig. 3h). In agreement with these findings, we found increased levels of the ribosome biogenesis-associated proteins nucleolin (Ncl) and rRNA methyltransferase fibrillarin (Fbl)^39,40^ (Extended Data Fig. 4e). These observations revealed a novel role for CDK8 in regulating oncogenesis by mediating global protein synthesis.

### CDK8 mediates stemness and differentiation in MYC-driven medulloblastoma

Emerging evidence has shown that dysregulated ribosome biogenesis may influence cancer stem cell differentiation pathways, impacting tumor progression and therapeutic responses^41,42^. Using single-cell RNA-seq of samples from seven patients, we identified a large population of undifferentiated progenitor-like cells exhibiting elevated expression of ribosomal genes in G3-MB (Fig. 4a,b and Extended Data Fig. 5a). CDK8 is predominantly expressed in undifferentiated populations and exhibits a significant overlap with the ribosomal gene population. This pattern was also observed in two additional Group 3 MB murine models: the cMyc-overexpressing and dominant-negative Trp53 GP3 mouse model and the Myc and Gfi1 co-expression GP3 mouse model, suggesting the role of CDK8 in mediating stemness and differentiation through the CDK8-mediated translation program in G3-MB (Fig. 4c). GSEA indicated that the loss of CDK8 resulted in strong negative enrichment in the transcriptomic profile associated with stemness gene sets and positive enrichment in differentiation gene sets (Fig. 4d and Extended Data Fig. 5b). Additionally, both genetic knockdown or pharmacological inhibition of CDK8 diminished the capacity for self-renewal in MB cultures, as assessed by aldehyde dehydrogenase activity, a marker associated with stem-like properties such as self-renewal and the capacity to differentiate into multiple cell types, and the efficacy of neurosphere formation, suggesting that CDK8 promotes self-renewal and stem-like phenotypes in MYC-driven MB (Fig. 4e,f and Extended Data Fig. 5c,d).

**Fig. 4.**
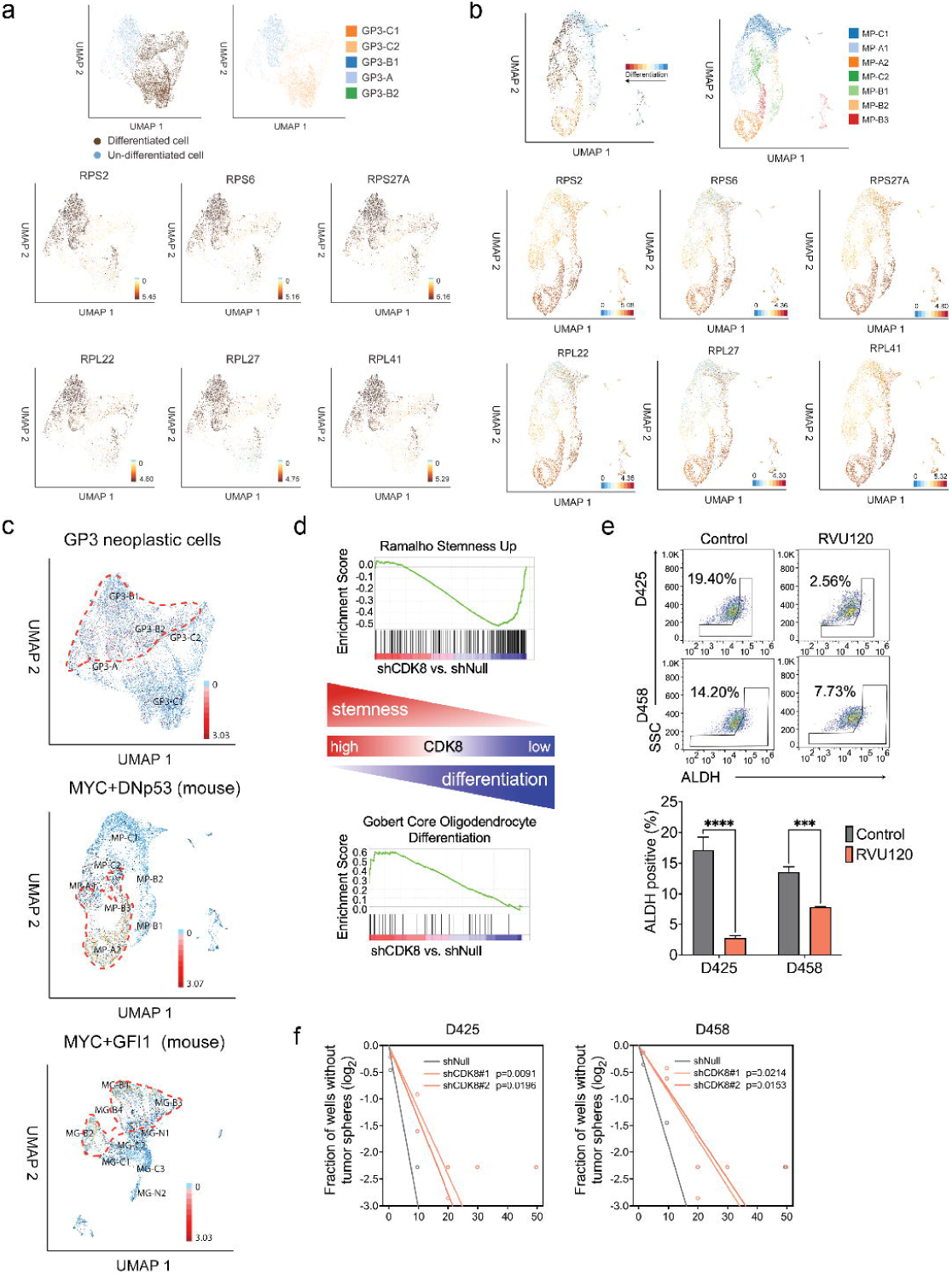
| CDK8 mediates stemness and differentiation in MYC-driven medulloblastoma. **a,** Harmony alignment of 12595 GP3 neoplastic cells from 7 MB patient samples demonstrates differentiated cells and undifferentiated cell populations (top). GP3 tumors formed 5 clusters that included 2 differentiated (GP3-C2, GP3-C1), 2 progenitor (GP3-B1, GP3-B2) and a mitotic (GP3-A) subpopulation. Representative ribosomal gene expression is shown (bottom). **b,** UMAP of 5422 cells of c-Myc and dominant-negative Trp53 GP3 mouse model demonstrates differentiated cell and undifferentiated cell populations (top). Representative ribosomal gene expression is shown (bottom). **c,** Harmony alignment or UMAP showing the expression of CDK8 in human patient cells and cells from mouse MB models. **d,** GSEA showing depletion of CDK8 reduced stemness gene sets and promoted differentiation gene sets in MB cells. **e,** Identification of the brain tumor-initiating cell fraction in MB cells by ALDH expression demonstrates a decrease in the ALDH+ fraction following IC50 RVU120 treatment for 48 hours. Mean ± SD. Statistical analysis: one-way ANOVA. **f,** Sphere formation efficiency and self-renewal capacity were measured using extreme *in vitro* limiting dilution assays (ELDA) in two MB cell lines. P values were determined using the likelihood ratio test.

### CDK8 transcriptionally regulates ribosomal genes

CDK8 plays a crucial role as part of the Mediator complex^19^. To determine whether CDK8 functions as a transcriptional activator affecting the translation program, we performed a genome-wide analysis to map the occupancy of CDK8 and key histone markers using CUT&RUN in three G3-MB cell lines. CDK8 binding peaks were identified in both the promoter and enhancer regions (Fig. 5a and Extended Data Fig. 6a). Next, we obtained gene annotations for CDK8 binding peaks and performed functional enrichment analysis to identify the predominant biological themes among these genes. We found that the pathways associated with translation were enriched in all three MB cell lines (Fig. 5b). Further analysis revealed that the predominant genes within these pathways were cytosolic and mitochondrial ribosomal genes, suggesting that CDK8 regulates ribosomal gene transcription (Fig. 5c).

**Fig. 5.**
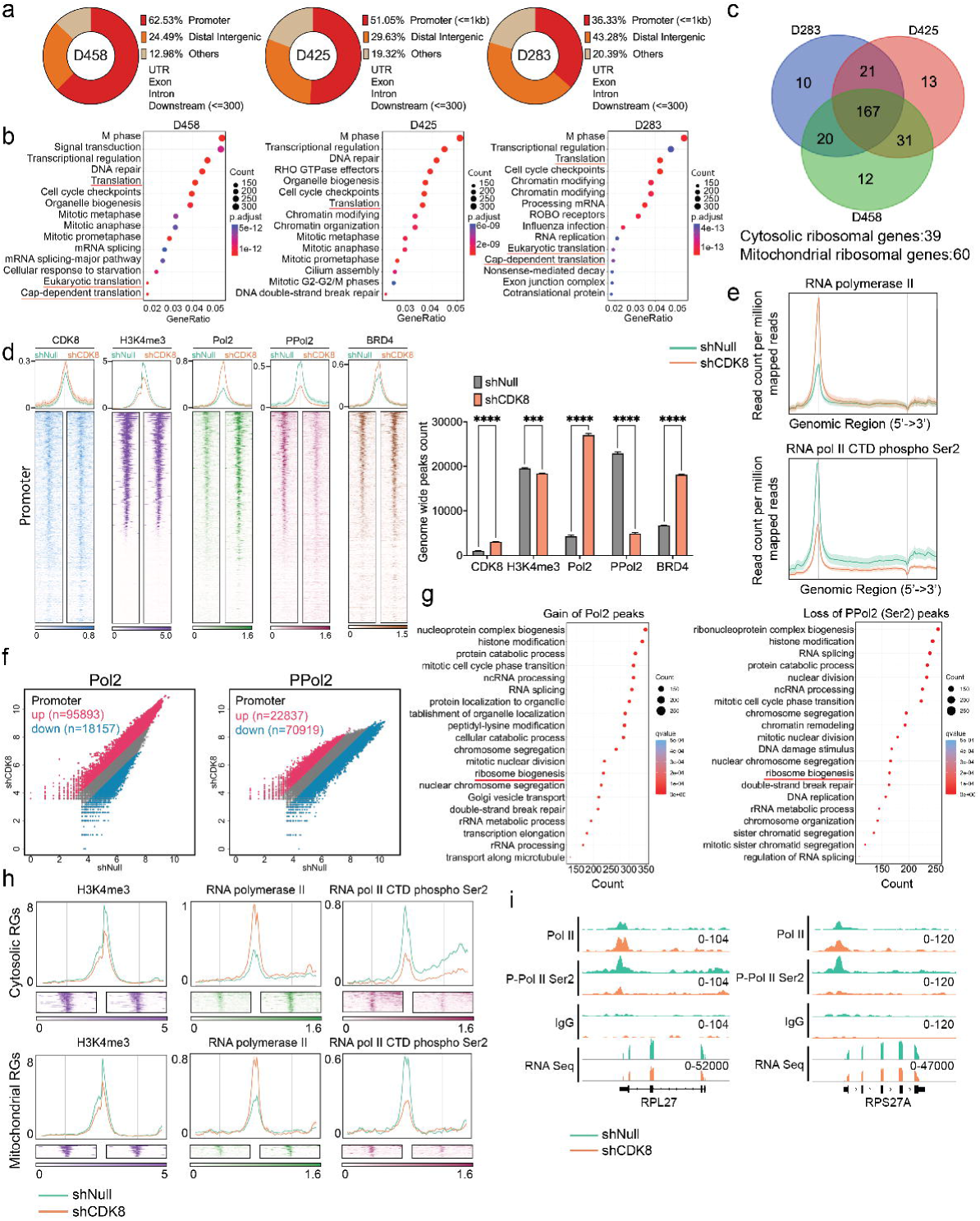
| CDK8 transcriptionally regulates ribosomal genes. **a**, Pie chart showing CDK8 is primarily localized at promoter and putative enhancer sites in G3-MB cell lines. **b,** Pathway enrichment analysis of CDK8 binding genes inferred from CUT&RUN profiling. Translation-related pathways are enriched in three G3-MB cell lines. **c,** Venn diagram showing overlapping of CDK8 binding genes associated with mRNA translation pathways in MB cell lines. The majority of these genes are related to cytosolic and mitochondrial ribosomal genes. **d,** Bar plots and heatmaps showing CUT&RUN peak numbers and signals of CDK8, H3K4me3, RNA Pol II, phosphor-RNA Pol II, and BRD4 in MB cell lines. The signals are displayed within a region spanning ± 3kb around the transcription start site (TSS). Mean ± SD. Statistical analysis: one-way ANOVA. **e,** Average distribution of RNA Pol II and phosphor-RNA Pol II peaks showing a significant increase in RNA Pol II and decrease in phosphor-RNA Pol II across the gene body following CDK8 knockdown. **f,** Scatter plot showing the dynamics of RNA Pol II and phosphor-RNA Pol II enrichment changes between shNull and shCDK8 conditions for the promoter regions. Peaks with at least 1.5-fold enrichment change upon CDK8 knockdown are highlighted (gain in red, loss in blue). The number of peaks is shown in the plots. **g,** Enrichment analysis identifies mRNA translation related pathways as significantly enriched among genes with an increase in RNA Pol II peaks or a decrease in phosphorylated phosphor-RNA Pol II following CDK8 knockdown. **h,** Average distribution and heatmaps of H3K4me3, RNA Pol II, and phosphor-RNA Pol II signals on cytosolic and mitochondrial ribosomal genes. **i,** Representative examples of RNA Pol II and phosphor-RNA Pol II binding sites on ribosomal genes observed following CDK8 knockdown.

Upon knockdown of CDK8, we found a significant decrease in its genome-wide occupancy, affecting genes associated with chromatin remodeling and translation pathways, indicating a role for CDK8 in the transcriptional regulation of translation program (Extended Data Fig. 6b,c). Next, we assessed the occupancy of RNA Pol II, phospho-Pol II (Ser2), and typical histone markers (H3K4me3, BRD4, H3K4me1, and H3K27me3). While CDK8 depletion minimally affected canonical enhancers, such as BRD4 and H3K4me1, as well as the repressive marker H3K27me3, it induced significant changes in chromatin dynamics, including RNA Pol II pausing and reductions in phospho-Pol II and H3K4me3 at promoter regions, which are crucial for gene activation and the initiation of transcription (Fig. 5d and Extended Data Fig. 6d). Knockdown of CDK8 caused RNA Pol II to primarily pause at the promoter regions, whereas the decrease in phosphor-Pol II continued from 5’ to 3’ across the gene body (Fig. 5e and Extended Data Fig. 6e). Among the peaks displaying at least a 1.5-fold change following CDK8 knockdown, there was an over five-fold increase in RNA Pol II-binding sites and a three-fold decrease in phospho-Pol II-binding sites (Fig. 5f). These findings suggest that CDK8 impacts the phosphorylation of RNA polymerase II, thereby influencing gene expression regulation and efficiency. Additionally, the differential alterations in RNA Pol II and phospho-Pol II peaks markedly contributed to various pathways, including rRNA metabolic processes and ribosome biogenesis, indicating that CDK8 regulates ribosomal gene expression (Fig. 5g). Similar chromatin alterations in RNA Pol II and phospho-Pol II were observed in both cytosolic and mitochondrial ribosomal genes following CDK8 knockdown (Fig. 5h). These chromatin changes were associated with ribosomal gene expression, as evidenced by the overlap peak track of CUT&RUN and RNA-Seq (Fig. 5i). Furthermore, treatment with RVU120 inhibited CDK8 activity, leading to decreased binding of Pol II to promoters and reduced occupancy of phospho-Pol II, supporting the finding that CDK8 modulates ribosomal gene expression (Extended Data Fig. 7a-d).

### Hyperactive ribosome biogenesis in MYC-driven Group 3 medulloblastoma

Aberrant protein synthesis is a common characteristic of MYC-driven cancers^43,44^. The mammalian target of rapamycin (mTOR) plays a key role in protein synthesis by regulating translational initiation, elongation, and ribosome biogenesis. Although mTOR inhibitors have been investigated in various types of cancer, their efficacy in MYC-driven MB remains unclear because of conflicting results from different studies^45,46^. Studies have reported that phospho-4EBP1 is significantly enriched in SHH-MB compared to G3-MB and that mTOR inhibition was not able to suppress G3-MB tumorigenesis^47,48^. In contrast, recent findings have demonstrated that the third-generation mTOR inhibitor RapaLink-1 enhances survival in a genetically engineered mouse model of MB (Glt1-tTA:TRE-MYCN/Luc mice)^49^. To address this paradox, we performed gene set variation analysis (GSVA) on gene expression data from 763 MB patient samples^50^. Our analysis revealed hyperactive mRNA translation and ribosome biogenesis in the G3-MB cells (Fig. 6a). Subsequently, multiplex IHC was performed on the G3-MB patient samples stained for CDK8, p-4EBP, p-S6, c-MYC, and RPS12. Consistent with our gene-level findings, the staining intensity of all these protein markers was significantly higher in G3-MB than in non-tumor control regions, suggesting that targeting protein synthesis could be a potential therapeutic strategy for G3-MB (Fig. 6b and Extended Data Fig. 8a). Next, we evaluated the efficacy of mTOR inhibition in an MB xenograft model. Using the second-generation mTOR inhibitor TAK-228, known for its ability to penetrate the blood-brain barrier, we observed a significant increase in survival and enhanced apoptosis in the treated cohort compared to the control group, indicating that targeting mTOR could be a therapeutic approach for MYC-driven MB (Fig. 6c-e).

**Fig. 6.**
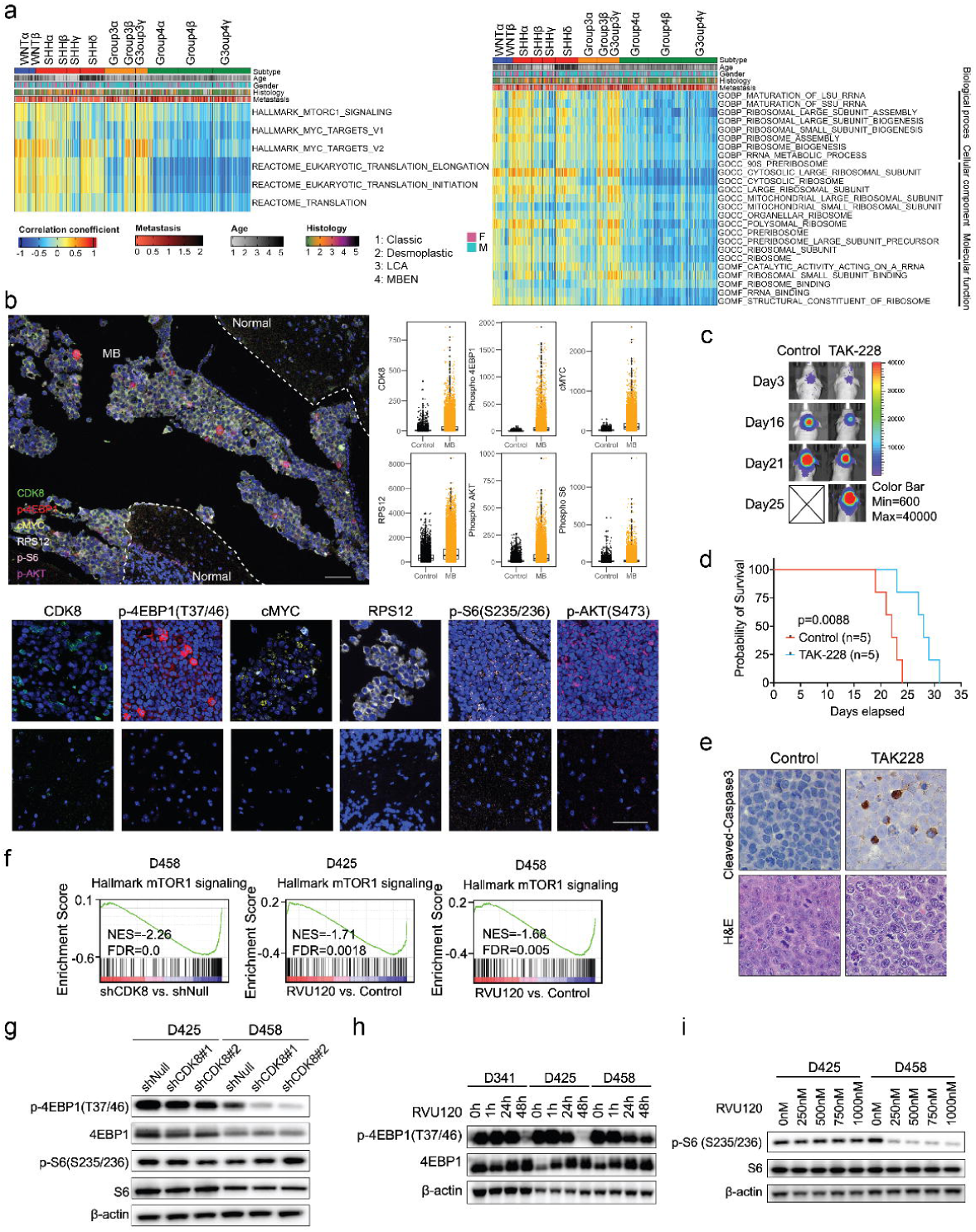
| Hyperactive ribosome biogenesis in MYC-driven Group 3 medulloblastoma. **a**, Gene set variation analysis of Cavalli’s patient samples (n=763) revealed that subtypes Group3β and 3γ were enriched with GO terms related to mRNA translation and ribosome biogenesis. **b,** Multiplex IHC on G3-MB patient samples using CDK8, p-4EBP1, c-MYC, RPS12, p-S6, and p-AKT antibodies. p<0.05 in all biomarker groups. Statistical analysis: unparied t-test. **c,** Representative bioluminescence images of mice treated with TAK-228 (1mg/kg, daily, oral gavage) compared with those of the control cohort. **d,** Kaplan–Meier survival curves for animals treated with control or TAK-228. Statistical analysis: Log-rank test. **e,** IHC analyses of cleaved caspase 3 protein expression in xenografts mice. Three mice from each group were sacrificed 18 days after tumor implantation. Original magnification, ×40. **f,** GSEA plots of representative gene sets involved in mTOR signaling following CDK8 depletion. Normalized enrichment score (NES) and false discovery rate (FDR) are indicated. **g,** Immunoblot showing the levels of p-4EBP1 and p-S6 following CDK8 knockdown. **h,** Immunoblot showing IC50 RVU120 treatment inhibited the levels of p-4EBP1 across three MB cell lines. **i,** Immunoblot showing the level of p-S6 upon treatment with various dose of RVU120.

GSEA of RNA-Seq data from genetic knockdown or pharmacological inhibition of CDK8 demonstrated significant downregulation of gene sets associated with mTOR signaling (Fig. 6f). To investigate the mechanism by which CDK8 functions in mTOR signaling, we assessed two major substrates of mTORC1: S6K1 and 4EBP1. Upon genetic knockdown of CDK8 or dose-dependent treatment with RVU120, MB cells showed decreased phospho-4EBP1 but not phosphor-S6 (Fig. 6g,h). Only D458 cells exhibited decrease phosphorylation of phosphor-S6 upon RVU120 treatment (Fig. 6i). This possible due to the distinct genetic backgrounds of the MB cells; D425 carries a loss-of-function V274C TP53 mutation while D458 has WT-TP53. Recent study has suggested that hematopoietic stem and progenitor cells exhibit stage-specific translational programs through CDK1-dependent mechanism, suggesting a potential crosstalk between CDKs and mTOR signaling^51^. To further examine the role of CDK8 in mTOR signaling, we examined the localization of CDK8 using two CDK8 antibodies across various MB cells, NHA, and NIH3T3 cells. Immunofluorescence analysis revealed predominant CDK8 expression within the nucleus, accompanied by additional expression in the cytoplasm (Extended Data Fig. 8b). Moreover, co-localization of CDK8 and phosphor-4EBP1 was observed (Extended Data Fig. 8c). Further validation through co-immunoprecipitation assay confirmed specific binding between CDK8 and phosphor-4EBP1 (Extended Data Fig. 8d). These findings suggest that CDK8 may regulate protein synthesis through interaction with 4EBP1, thus activating eIF4E-cap-dependent translation. Therefore, concurrent modulation of the CDK8 and mTOR pathways could potentially synergize to enhance therapeutic outcomes in MYC-driven MB.

### Synergistic Targeting of CDK8 and mTOR for MYC-Driven Medulloblastoma

Given the similar impact of mTOR and CDK8 inhibitors on the suppression of protein synthesis in MB cells, we examined whether the simultaneous inhibition of CDK8 and mTOR could synergistically impede the growth of MB cells. CDK8 knockdown cells showed reduced sensitivity to Torin1, an ATP-competitive inhibitor that blocks mTORC1 and mTORC2, as demonstrated by the lower IC50 compared to control cells (Extended Data Fig. 8a). Next, we performed a combination study using increasing doses of RVU120 and the mTOR inhibitor Torin 1 on MB cells. Dual inhibition resulted in a significant synergistic effect on the lethality and proliferation of MB cells (Fig. 7a,b). Subsequent evaluation using the Chou-Talalay method and Bliss synergy model confirmed this synergistic effect (Fig. 7c,d and Extended Data Fig. 8b). Flow cytometry analysis revealed that combination treatment enhanced the apoptosis of MB cells (Fig. 7e and Extended Data Fig. 8c). Consistent with these results, dual inhibition significantly decreased the levels of p-4EBP1 and p-S6 and reduced p-STAT1 and phospho-Pol II levels, further emphasizing CDK8’s role in governing both protein synthesis and chromatin dynamics (Fig. 7f).

**Fig. 7.**
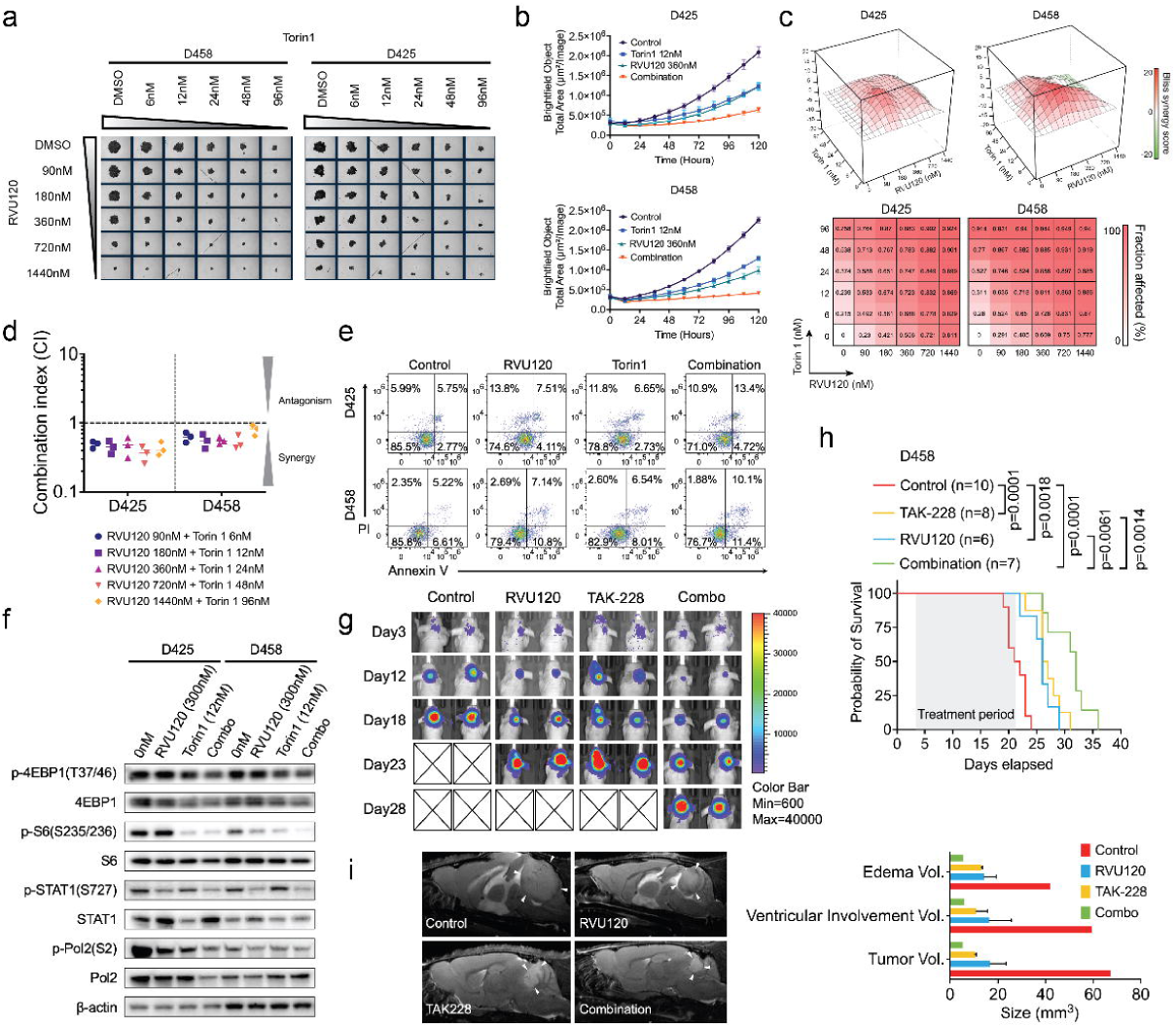
| Synergistic targeting of CDK8 and mTOR in MYC-Driven Medulloblastoma. **a** Dose-dependent assay of the combined treatment with RVU120 and Torin1 on Day 5 in D425 and D458 cells. **b,** Real-time proliferation assay quantifying the combined treatment with RVU120 and Torin1 using the Incucyte Live-Cell Analysis System. **c,** Heatmap representation of the Fraction Affected (FA) and the Bliss interaction index across the five-point dose range of RVU120 and Torin1 in D425 and D458 cells. Mean values of triple biological experiments are shown. **d,** The combination index of RVU120 and Torin1 using chou-talalay method. The mean combination index (horizontal bar) was determined from three independent experiments using D425 and D458 cells. The results for each experiment are indicated by symbols. **e,** Apoptosis assay following combined treatment with RVU120 and Torin1. D425 and D458 cells were treated for 48 h before staining with PI and Annexin V. **f,** Effects of the combination of RVU120 and Torin1 on protein synthesis markers, phospho-Pol2 and phospho-STAT1, in D425 and D458 cells after 48 h of treatment. **g,** The nude mice injected with D458 cells were treated with vehicle, RVU120 (40 mg/kg), TAK-228 (1 mg/kg), or their combination. Representative bioluminescence images are shown. Color scales indicate bioluminescence radiance in photons/sec/cm2/steradian. **h,** Kaplan-Meier survival curve of D458 xenograft mice. The treatment period is indicated by shaded boxes. Statistical analysis: log-rank (Mantel-Cox) test. **i,** Representative Sagittal T2-weighted turboRARE MRI of D458 xenografted mice at 22 days. White arrows indicate tumors. MRI volumetric analysis (right) demonstrates trend toward decreased tumor volume in each cohort.

To explore synthetic lethality *in vivo*, we assessed the efficacy of RVU120 and TAK-228 administered individually or in combination in D458 xenograft mouse model. Initially, the IVIS signals indicated similar tumor sizes in all groups. Tumor growth notably decelerated in the treated mice, particularly in the cohort that received combination treatment (Fig. 7g). Consistent with the *in vitro* data, mice receiving the combination treatment showed the most effective therapeutic outcomes, characterized by prolonged overall survival and reduced tumor burden, as determined by MRI (Fig. 7h,i). Additionally, hematological analyses conducted prior to sacrificing the mice revealed that administration in each group did not induce notable acute hematological toxicity, as evidenced by stable white blood cells, neutrophils, and lymphocytes (Extended Data Fig. 8d-f). Taken together, these studies established the therapeutic efficacy of combination treatment with mTOR and CDK8 inhibitors *in vivo* and *in vitro*, opening an alternate path for biologically based therapeutic trials in MYC-driven MB.

## Discussion

Medulloblastoma is the most common and lethal pediatric brain tumor^1,52–54^. It is crucial to identify disease vulnerabilities and develop therapies that target specific mechanisms. Here, we identified MYC-driven medulloblastoma as one of the most significantly affected cancer types following CDK8 depletion, demonstrating the essential role of CDK8 in driving medulloblastoma growth, and unveiling its previously unexplored oncogenic significance in MB. Our data expand on previous studies that identified a link between CDK8 and MYC and provides a new mechanism by which CDK8 may facilitate MYC driven tumorigenesis^17,55,56^.

In G3-MBs, approximately 17% of cases demonstrate high-level MYC amplification, a defining characteristic contributing to widespread treatment failure in children diagnosed with MYC-amplified MB despite current therapies^57^. MYC, which functions as a pleiotropic transcription factor, promotes the proliferation of neural progenitor cells in malignant stem cells by modulating overall gene expression and regulating critical cellular processes^58^. Although MYC can drive cerebellar stem cell proliferation *in vitro*, it is insufficient to maintain long-term growth in animal models. A previous study revealed that cerebellar stem cells require both MYC overexpression and mutant Trp53 to generate aggressive MB upon orthotopic transplantation^59^. Similar studies have demonstrated that the combination of MYC with GFI1 or MYC with SOX2 leads to rapid formation of highly aggressive cerebellar tumors using stem cells or astrocyte progenitors^60,61^. Given the significant role of CDK8 in G3-MB identified in our study, it is likely to collaborate with MYC to promote a stem cell-like state and hinder MB cell differentiation. It will be of great interest to determine in future studies whether CDK8 and MYC overexpression in cerebellar stem cells is sufficient to drive tumorigenesis and form G3-MB tumors.

Dysregulation of protein synthesis is a common characteristic of MYC-driven cancers and is marked by increased Pol I-mediated ribosomal rDNA transcription and mTOR/eIF4E-driven mRNA translation^12,16^. Previous studies have demonstrated the robust efficacy of PI3K/mTOR inhibitors in inhibiting the growth of MB cells derived from MYC+DNp53 transfected stem cells, both *in vitro* and *in vivo*^62^. Our findings demonstrated that G3-MB exhibited a dysregulated protein synthesis profile, predominantly comprising undifferentiated progenitor-like cells with significantly elevated expression of ribosomal genes, indicating vulnerability for considering protein synthesis as a potential target for treatment. CDK8 depletion remarkably repressed pathways associated with ribosome biogenesis and mRNA translation. Subsequently, we investigated the mechanisms through which CDK8 regulates these cellular activities. Our findings reveal that CDK8 plays a dual role in the regulation of protein synthesis. First, as a cyclin-dependent kinase, CDK8 interacts with and contributes to the phosphorylation of 4EBP1, a crucial downstream target of mTOR signaling. This phosphorylation by CDK8 potentially leads to the dissociation of 4EBP1 from EIF4E, allowing the assembly of the translation initiation complex, consequently facilitating eIF4E-dependent translation. Second, as a dissociable part of the mediator complex, CDK8 inhibition results in decreased phosphorylation of RNA Pol II, consequently affecting the targeted suppression of gene expression, specifically of genes linked to ribosomal function. A previous study established a correlation between CDK8 levels and the mTOR pathway in acute lymphoblastic leukemia, suggesting that CDK8 regulates protein synthesis not only within a subset of MB, but also in other types of cancer^63^.

Despite the importance of CDK8 in regulating protein synthesis in MB, there remains unclarity regarding how dysregulation of protein synthesis contributes to the development and progression of cancer. One possibility is that dysregulated translation promotes cell growth, proliferation, and metastasis^64^. This is supported by the observation that cancer cells frequently develop a strong addiction to protein synthesis to adapt to different microenvironments, providing vulnerability that can be effectively targeted by inhibiting protein synthesis in these cancer types^65^. Another possibility is that changes in translational dysregulation may affect specific molecular or cellular processes that contribute to cancer initiation and progression^66,67^. Studies have demonstrated that aberrant protein synthesis leads to changes in the expression of specific genes by affecting chromatin dynamics via epigenetic mechanisms^68,69^. These findings align with the recognized role of CDK8 in the Mediator complex, suggesting that CDK8 cooperates with MYC or other transcription factors to modulate transcriptional regulation, chromatin modifications, and the overall chromatin landscape, thereby impacting gene expression and crucial cellular processes essential for development, stability, and disease states, such as cancer.

We demonstrated a novel therapeutic strategy for targeting MYC-driven MB using RVU120, a new specific and selective inhibitor of CDK8^31^. RVU120 exhibits sufficient pharmacological properties, such as high oral bioavailability and brain penetration. A Phase 1 trial in patient with AML or high-risk MDS patients showed good tolerability with acceptable toxicity and signs of clinical activity (NCT04021368). New Phase 2 studies are currently underway. Despite the promising results obtained with CDK8 inhibitors in our preclinical study, some challenges remain to be addressed. One challenge is to identify patient populations that will benefit the most from CDK8 inhibitor therapy. Another challenge is the development of strategies to overcome the potential resistance mechanisms that may arise during treatment. In conclusion, these data suggest that CDK8 inhibitors are promising agents for MYC-driven medulloblastoma therapy and provide a mechanistic basis for future research.

## Methods

### Cell lines

The medulloblastoma cell line D425 was purchased from Millipore Sigma (SCC290). D458 was purchased from Cellosaurus (CVCL_1161), D283 from ATCC (HTB-185), and D341 from ATCC (HTB-187). MB002 was provided by Dr. Martine Roussel (St. Jude Children’s Research Hospital). HDMB03 was provided by Dr. Mahapatra of (University of Nebraska). Human astrocytes were cultured in complete Astrocyte Medium (ScienCell, 1801). MAF1433 cells were isolated and cultured from the primary tumor of a Group 3 MB patient. The D425 and D458 cell lines were cultured in DMEM, 10% FBS, 1% 1× penicillin/streptomycin solution, 1% 1× L-glutamine, and 1% sodium pyruvate. D283 cells were cultured in DMEM (Thermo Fisher) supplemented with 10% FBS, 1 mM sodium pyruvate, 1× penicillin/streptomycin solution (Cellgro), and 1× non-essential amino acids (Millipore Sigma). HDMB03 cells were cultured in 90% RPMI 1640, 10% FBS, and 1× penicillin/streptomycin. D341 and MB002 were cultured in neurobasal medium (Sigma, SCM003) containing 2% B-27, 1 μg/ml heparin, 2 mM L-glutamine, 1% penicillin/streptomycin, 25 ng/ml fibroblast growth factor (FGF), and 25 ng/ml epidermal growth factor (EGF). All cell lines were cultured at 37°C in 95% air and 5% CO_2_. All cell lines tested negative for Mycoplasma. Cell proliferation assays and live-cell imaging were performed using the Incucyte SX5 Live-Cell Analysis System (Sartorius).

### Protein synthesis assay

The MB cells were plated at a density of 2,000 cells/well in a 96-well plate and cultured overnight. The next day, the cells were treated with either vehicle or RVU120 for 1, 24, or 48 h. The cells were then collected and centrifuged at 400 × g and resuspended in OPP (O-propargyl-puromycin) working solution (Cayman Chemical, 601100). The mixed cells were incubated for 30 min at 37°C for OPP labeling of translated peptides. Following incubation, cells were fixed, washed, and analyzed using flow cytometry.

### Transfection

shRNA vectors targeting CDK8 mRNA (#TRCN0000350344, # TRCN0000382350) and a non-targeting shRNA (control) were purchased from the Functional Genomics Facility at the University of Colorado Anschutz Medical Campus. Transfection was performed using the Lipofectamine 3000 Transfection Reagent (Invitrogen).

### Extreme limiting dilution assay

The cells were treated with the indicated concentrations of RVU120 and then seeded into 96-well ultra-low-attachment plates in neurosphere media at increasing concentrations from 1 to 250 cells/well. Cells were seeded from n = 5 wells (250 cells/well, 100 cells/well), n = 10 wells (10–50 cells/well), or n = 30 wells (1 cell/well) per condition. The cells were allowed to grow for 14 days, and the number of wells containing neurospheres was counted under a microscope.

### Aldehyde dehydrogenase assay

ALDH activity was measured using an Aldefluor kit (Stem Cell Technologies), according to the manufacturer’s instructions. Briefly, 1 × 10^5^ cells were resuspended in 0.5 mL Aldefluor buffer, separated equally into two tubes, and 5 μl of DEAB reagent was added to one tube as a negative control. Then 1.25 μl of Aldefluor Reagent was added to each tube and mixed well. After incubation at 37°C for 45 min and centrifugation, cells were stained with propidium iodide and then analyzed using a FlowSight Imaging Flow Cytometer (EMD Millipore).

### Neurosphere assay

Medulloblastoma cells were grown for 14 days in neurosphere medium. The spheres were disassociated and replanted in 100-, 10-, and single-cell suspensions on day 14. The cells were grown for an additional 14 d with or without RVU120. The spheres were imaged using the Incucyte S3 Live Cell Imaging System (Sartorius).

### Methylcellulose assay

2000 cells/3 mL were plated in a 1:1 mixture of 2.6% methylcellulose and complete growth medium. The cells were allowed to grow for two weeks. colonies were stained with nitrotetrazolium blue chloride (Sigma) at 1.5mg/mL in PBS for 24 h at 37°C and counted.

### Compounds

The CDK8 inhibitors Torin1 and TAK-228 were purchased from MedChemExpress, and RVU120 for animal studies was provided by Ryvu Therapeutics. The drugs were reconstituted in dimethyl sulfoxide (DMSO). An equivalent amount of DMSO at the highest concentration of the drug was used for each experiment as a vehicle control.

### Drug interaction assay

Medulloblastoma cells were plated in 96-well low-attachment plates and subjected to dose-response assessments for individual drugs, as well as various concentrations of drug combinations, with DMSO (0.1%) and media serving as controls. The growth inhibition was quantified using the CellTiter 96 AQueous Non-Radioactive Cell Proliferation Assay (Promega) and the Incucyte SX5 Live-Cell Analysis System (Sartorius). At least five independent trials were conducted to ensure reproducibility of the results. The Chou-Talalay median-effect model and the Bliss independence dose-response surface model were used to classify whether the two drugs interacted in an antagonistic, additive, or synergistic manner. For the Chou-Talalay median-effect model, CI > 1 indicated antagonism, CI = 1 demonstrated activity, and CI < 1 indicated synergistic interactions.

### Animal studies

Female athymic Nude *Foxn1*^nu^ and female NOD scid (NSG, #5557) gamma mice aged 4–8 weeks were used for orthotopic xenograft studies. D458 cells were collected and resuspended in a single cell suspension of 20,000 cells/3 μl in non-FBS medium. The mice were monitored daily for tumor growth and euthanized when 15% weight loss was reached. To monitor tumor growth in D458 xenograft mice, the mice were injected intraperitoneally with 10 μl/g of 15 mg/mL D-luciferin potassium salt solution (Gold Biotechnology) and imaged using the Xenogen IVIS 200 In Vivo Imaging System (PerkinElmer). Tumor bioluminescence was analyzed using the Living Image 2.60.1 software (PerkinElmer). The mice used in this study were kept in a sterile envrioment under 12/12-h light/dark cycle, 21-23 °C and 40–60% humidity at University of Colorado, Anschutz Medial Campus, Aurora, USA.

The mice were administered a daily dose of 40 mg/kg RVU120 or 1 mg/kg TAK-228 via oral gavage. RVU120 was dissolved in water and TAK-228 was prepared by dilution in N-methyl-2-pyrrolidone (NMP) and subsequent suspension in a 15% polyvinylpyrrolidone solution for administration. In the combination treatment group, mice received RVU120 initially, followed by a 2-hour intermission before the administration of TAK-228. All mice were treated with the respective drugs 2-4 hours before sacrifice, and blood was extracted for hematological toxicity analysis.

### Magnetic resonance imaging

For *in vivo* MRI acquisitions, the mice were anesthetized shortly before and during the MR session using a 1.5% isoflurane/oxygen mixture. The anesthetized mice were placed on a temperature-controlled mouse bed below a mouse head array coil and inserted into a Bruker 9.4 Tesla BioSpec MR scanner (Bruker Medical). First, T2-weighted turboRARE images were acquired using the following parameters: repetition time (TR), 3268 ms; echo time (TE), 60 ms; RARE factor, 12 and 8 averages; FOV, 20 mm; matrix size, 350 × 350; slice thickness, 700 µm; 24 sagittal and axial slices; in-plane spatial resolution, 51 µm. A diffusion-weighted EPI sequence with 6 b values was then used using four axial slices covering all tumor lesions and unaffected brain tissue. Tumor regions were manually segmented on T2-weighted images by placing hand-drawn regions of interest (ROI) and the volume was calculated in mm3. The apparent diffusion coefficient (ADC; s/mm2) was calculated using diffusion-weighted imaging maps as a criterion for tumor cellularity. All acquisitions and image analyses were performed using the Bruker ParaVision NEO software (Bruker Medical).

### Immunofluorescence

The cells were washed and seeded onto polylysine-coated slides, and then fixed with 4% paraformaldehyde for 15 min at room temperature, permeabilized with 0.2% Triton X-100 in PBS for 15 min, and incubated in 3% BSA diluted in 0.05% Triton X-100 for 30 min at room temperature on a shaker. After blocking, the cells were incubated with the primary antibodies. The following antibodies were used: Phospho-4EBP1 (Santa Cruz Biotechnology, sc-293124, 1:50), CDK8 (Santa Cruz Biotechnology, sc-13155, 1:50), CDK8 (Abcam, ab224828, 1:200), CDK8 (Invitrogen, PA5-11500, 1:200), fibrillarin (Abcam, EPR10823, 1:200), nucleolin (Abcam, EPR7952, 1:200), and ribosomal RNA antibody Y10B (Abcam, ab171119, 1:200) for 1 h at room temperature. After washing with 0.05% Triton X-100, cells were incubated with Alexa Fluor 647-or Alexa Fluor 488 conjugated secondary antibody (1:500) for 1 h at room temperature in the dark, washed with PBS, and mounted using ProLong Gold antifade reagent containing DAPI (Sigma). Images were acquired using an inverted epifluorescence microscope at × magnification of 40x.

### Flow cytometry

Cells were fixed with 4% formaldehyde for 15 min at room temperature. The fixed cells were washed and permeabilized with methanol on ice for 10 min. Cells were stained with active caspase-3 (BD Biosciences). The apoptosis was measured by eBioscienc Annexin V apoptosis detection kit FITC (Thermo Fisher). Flow cytometry analysis was performed using an Amnis FlowSight flow cytometer (Millipore).

### Western blotting

Western blotting was performed as described previously^23^. Antibodies used for western blot analysis were from the following sources: β-actin (Cell Signaling, 8457, 1:2000), CDK8 (Cell Signaling, 4101, 1:1000), 4EBP1 (Cell Signaling, 9644S, 1:1000), Phosphor-4EBP1 (Cell Signaling, 2855S, 1:1000), STAT1 (Cell Signaling, 9176S, 1:1000), Phosphor-STAT1 (Cell Signaling, 8826S, 1:1000), S6 (Cell Signaling, 2217T, 1:1000), Phosphor-S6 (Cell Signaling, 4858T, 1:1000), Pol II (Cell Signaling, 2629S, 1:1000), and Phosphor-Pol II-Ser2 (Cell Signaling, 13499, 1:1000).

### Co-immunoprecipitation

Co-IP was performed using a Universal Magnetic Co-IP kit (Active Motif). Briefly, 200ug nuclear or whole-cell extract protein was mixed with 5 ug CDK8 antibody in a final volume of 500 μl Complete Co-IP buffer in a pre-chilled tube. Incubate The mix for 1-4 hours at 4 °C on a rotator. next day, add 25 μl protein G Magnetic Beads to each reaction and incubate 1hour at 4 °C on a rotator. The immunocomplexes were then pelleted, washed four times at 4 °C, and subjected to SDS-PAGE and western blotting using anti-phospho-4EBP1 and anti-4EBP1 antibodies.

### Immunohistochemistry

For histological analysis, tumors from experimental mice were dissected and either frozen or preserved in 10% formalin. The samples were rinsed with PBS, fixed in 4% paraformaldehyde overnight at 4°C, and embedded in paraffin. Antigen retrieval was performed by the application of citrate buffer pH 6.00 for 20 min. Slides were then incubated with caspase 3 Cell Signaling Technology, #9662) or PLK1 (Abcam, ab109777) antibody and H&E overnight at 4 °C. The secondary antibody conjugated to horseradish peroxidase was detected using the Dako EnVision Kit for 3,3′-diaminobenzidine.

### RNA sequencing

RNA was isolated from cells under the indicated experimental conditions using a Qiagen miRNAeasy kit (Valencia) and measured using an Agilent Bioanalyzer (Agilent Technologies). Illumina Novaseq 6000 libraries were prepared and sequenced by Novogene (CA, USA) or the Genomics and Microarray Core Facility at the University of Colorado Anschutz Medical Campus. High-quality base calls at Q30 ≥ 80% were obtained with approximately 40 M paired paired-end reads. Sequenced 150bp pair-end reads were mapped to the human genome (GRCh38) by STAR 2.4.0.1, read counts were calculated by R Bioconductor package GenomicAlignments 1.18.1, and differential expression was analyzed with DESeq2 1.22.2 in R. Further analysis by GSEA was performed using GSEA v2.1.0 software with 1,000 data permutations and Cytoscape v3.10.1.

### CUT&RUN

A total of 500,000 cells per reaction were harvested and captured using 10 µL of pre-activated ConA beads (EpiCypher). Beads with attached cells were incubated at room temperature for 10 min to ensure complete adsorption. Subsequently, 50 µL of cold antibody specific to the reaction was added to each sample. The antibodies used for cu and run were CDK8 (Cell signaling, 4101S, 1:50), MYC (Cell signaling, 13987S, 1:50), Pol II (Cell signaling, 2629S,1:50), phospho-Pol II (cell signaling, 13499S, 1:50), H3K4ME1 (Abcam, ab8895, 1:50), H3K4ME3 (EpiCypher, 13-0041K, 0.5 mg/ml), BRD4 (Cell signaling, 13440S, 1:50), H3K27ac (Active motif, 39133, 1:25), H3K27me3 (Cell signaling, 9733s, 1:50), and IgG (EpiCypher, 13-0042K, 0.5 mg/ml). The cells were then incubated overnight on a nutator at 4 °C and permeabilized using a buffer containing 5% digitonin. Next, 2.5 µL/reaction pAG-MNase (Epicypher) was added to each sample. The beads were gently resuspended by vortexing or pipetting to evenly distribute the enzymes. The mixture was incubated for 10 min at room temperature. Calcium Chloride (100 mM, 1 µL/reaction) was added to the reaction, followed by a 2-hour incubation at 4 °C. After incubation, 34 µL of Stop Master Mix was added to each tube, followed by a 10-minute incubation at 37 °C. The tubes were then quick-spun and placed on a magnet for slurry separation, and the clear supernatants were transferred to 8-strip tubes for DNA purification. Libraries were prepared using the NEBNext Ultra II DNA Library Prep kit and sequenced using NovaSeq PE150.

CUT&RUN-seq reads were aligned to the reference human genome v.hg38 using BOWTIE v.2.3.4.1. Aligned reads were stripped of duplicate reads using Sambamba v.0.6.8. Peaks were called using the program MACS v.2.1.2, with the narrow peak mode using matched input controls and a q-value of 0.00001. Peaks in the blacklisted genomic regions identified by the ENCODE consortium were excluded using bedtools. For downstream analysis and visualization, bamCoverage was used to generate bigwig files and density maps were produced using IGV tools. Group 3 medulloblastoma enhancers were defined based on H3K27ac signals. Regions within 1kb of RefSeq transcription start site (TSS) locations and peaks with strong H3K4me3 signals typical of active promoters were subtracted from these signals. Annotation and visualization of the peaks were conducted using ChIPseeker v3.18. Differentially marked genes were calculated using DiffBind and DESeq2, based on the threshold of FDR < 0.05 and fold-change ≥ 2.

### Multispectral IHC

Tumor tissues were fixed in formalin and paraffin-embedded for multispectral imaging using the Vectra 3.0 Automated Quantitative Pathology Imaging System (Perkin Elmer). Four-micron sections mounted on glass slides were sequentially stained for human CDK8 (Abcam, ab224828), MYC (Abcam, ab168727), RPS12 (Abcam, ab167428), p-4EBP1-T37/46 (Abcam, ab75831), p-S6-S235/236 (Cell Signaling, 2211S), p-AKT-S473 (Leica, NCL-L-AKT-PHOS), and DAPI using a Bond RX autostainer (Leica). Slides were dewaxed (Leica), heat-treated in ER2 (epitope retrieval solution 2) antigen retrieval buffer for 20 min at 93 °C (Leica), blocked in antibody (Ab) Diluent (Perkin Elmer), incubated for 30 min with the primary antibody, 10 min with horseradish peroxidase-conjugated secondary polymer (anti-mouse/anti-rabbit, Perkin Elmer), and 10 min with horseradish peroxidase-reactive OPAL fluorescent reagents (Perkin Elmer). Slides were washed between staining steps with Bond Wash (Leica) and stripped between each round of staining by heat treatment in antigen retrieval buffer. After the final round of staining, the slides were heat-treated in ER1 antigen retrieval buffer, stained with spectral 4′,6-diamidino-2-phenylindole (Perkin Elmer), and coverslipped with ProLong Diamond mounting media (Thermo Fisher). Whole slide scans were collected using a 10× objective lens at a resolution of 1.0 μm. Approximately 30 regions of interest were selected from the tumor in areas near the tumor border or in the center of the tumor. Regions of interest were scanned for multispectral imaging with a 20× objective lens at a resolution of 0.5 μm. Multispectral images were analyzed using inForm software version 2.3 (Perkin Elmer) to unmix adjacent fluorochromes, subtract autofluorescence, segment the tissue into tumor regions and stroma, segment the cells into nuclear, cytoplasmic, and membrane compartments, and phenotype the cells according to cell marker expression.

### Microarray preparation and data processing

RNA from all surgical specimens was extracted, amplified, labeled, and hybridized to Affymetrix HG-U113 plus two microarray chips (Affymetrix). The scanned microarray data were background-corrected and normalized using the RMA algorithm, resulting in log 2 gene expression values. For the public microarray data, raw CEL files were downloaded from the Gene Expression Omnibus under accession number GSE85217 and normalized using the RMA algorithm. The gene expression array data generated using the Affymetrix Gene 1.1 ST array and U133 Plus 2.0, array platforms were merged to generate a combined value. For each platform, the contrast value per gene was calculated by subtracting the mean expression of that gene across all samples hybridized on that platform from each individual, and the resulting contrast values of the two platforms were then combined.

### Gene set enrichment analysis

Gene sets from MSigDB were downloaded and used to estimate biological activity. The ssGSEA algorithm in the R package GSVA (v.1.40.1) was applied to estimate signature enrichment in the bulk transcript datasets. The enrichment results of GO and pathways among differentially expressed genes were generated using the R package clusterProfiler (v.4.7.1).

### Single-Cell RNA-Seq

Single-cell RNA sequencing data were aligned against a composite reference consisting of mm10 and hg38 genomes to delineate transcripts originating from murine and human cancer cells using the Cell Ranger toolkit (version 4.0.0). The classification of cells as either human or murine was based on a threshold of 90% genome-specific reads. Cells falling below this threshold were identified as human-mouse chimeric multiplets and excluded from further analysis. Gene-barcode count matrices obtained from scRNA-seq were processed using the Seurat package (version 4.0.3) in R. Cells with fewer than 500 or more than 8,000 genes were excluded to eliminate low-quality samples and potential doublets. Cells with over 10% reads mapped to mitochondrial genes were filtered out. Log-normalization was applied to the filtered datasets, followed by principal component analysis to reduce the dimensionality. Utilizing Seurat’s elbow plot function, the top 25 principal components were selected for UMAP plot generation. Cell clusters were discerned via k-nearest neighbor unsupervised clustering and the resolution parameter was set to 1.2. Established markers from literature were used to annotate each cluster with its corresponding biological cell type.

### Survival analysis

R2: The Genomics Analysis and Visualization Platform (https://hgserver1.amc.nl/cgi-bin/r2/main.cgi?open_page=login) was used to delineate the association between gene expression levels and overall survival in patient samples. The log-rank P values, and Kaplan– Meier curves were calculated and plotted using the R package survival (v.3.2-11) and Prism GraphPad (v.10.0.2).

### Statistics analysis

The detailed statistic method are described in each individual figure legend. R package survival (v.3.2-11) and Prism GraphPad (v.10.0.2) were used for the statistics.

### Study approval

All patients provided written informed consent for molecular studies of their tumors, and the study protocol was approved by the ethics committee of the University of Colorado and Children’s Hospital Colorado (COMIRBs #95–500). All animal procedures were performed in accordance with the National Research Council’s Guide for the Care and Use of Laboratory Animals and approved by the University of Colorado, Anschutz Campus Institutional Animal Care and Use Committee.

## Acknowledgements

The authors appreciate the contribution made by the University of Colorado Tissue Histology Shared Resource, the Genomics Core and the small Animal Imaging Core, all supported in part by the Cancer Center Support Grant (P30CA046934). The authors thank Dr. Natalie Serkova for assistance with MRI imaging. The authors thank Dr. Craig Forester and Dr. Dylan Taatjes for review of the manuscript and valuable insight and discussion. The study was supported by the research grant from Morgan Adams Foundation (DW, RV), ACS IRG #16-184-56 from the American Cancer Society to the University of Colorado Cancer Center (DW), the Cancer League of Colorado, Inc. (DW, RV) and NIH grant RO1NS091219 (RV).

## Author Contributions

DW, CR and RV contributed to the design and implementation of the research. DW, BV, SV, ND, AP, YL and BB contributed to development and implementation of methodology. DW performed analysis of the results. KK and MM performed the blood brain barrier analysis for RVU120. DW, MM, TR and RV wrote and edited the manuscript. RV conceived the original project and supervised the project.

Conflict of Interest declaration: DW, CR, BV, SV, ND, AP, BB, YL, and RV report NO affiliations with or involvement in any organization or entity with any financial interest in the subject matter or materials discussed in this manuscript. KK, MM and TR are employees of RVYU therapeutics.

---

**Extended Data Fig.1.**
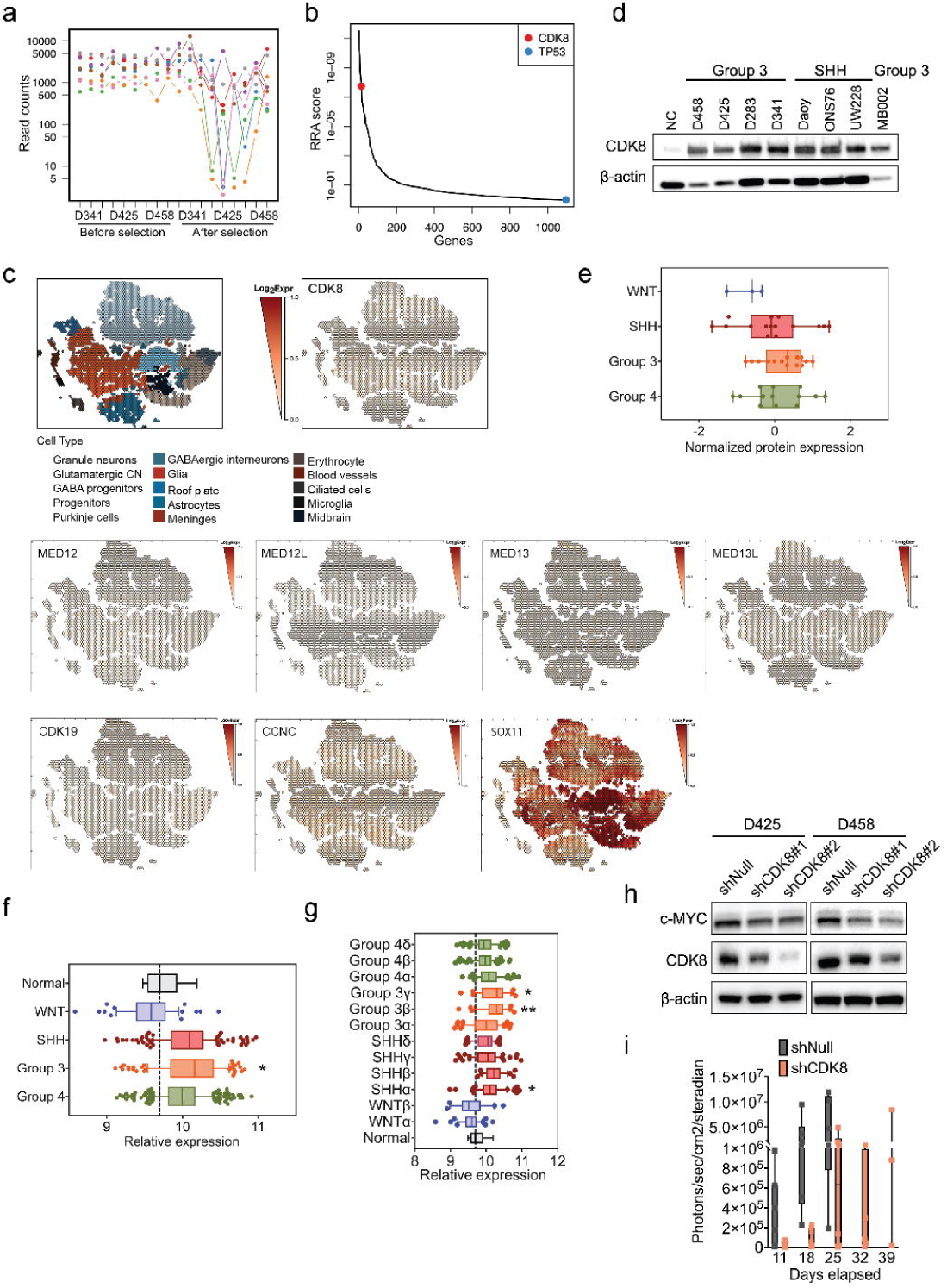
| Targeting CDK8 suppresses medulloblastoma progression. **a**, The read counts of sgRNAs targeting CDK8 in CRISPR-Cas9 screening across three MB cell lines. The columns are cell lines. Each dot represents a sgRNA targeting CDK8. **b,** The distribution of negative robust rank aggregation score in CRISPR-Cas9 screening. **c,** Scatterhex plots showing expression of CDK8 and Mediator complex related genes obtained from https://cellseek.stjude.org/cerebellum/. The gene expression of a hexgrid is summarized as the mean expression of all component cells and compared with predicted cell type mapped onto the scatterhex grid. SOX11, which was found to have higher expression in human fetal cerebellum and MB, was used as a control. **d,** Immunoblot showing the protein levels of CDK8 in Group 3, SHH human cell lines, and normal cerebellar human sample. **e,** Proteomic analysis demonstrating CDK8 expression across 4 subgroups of 45 MB patient samples. **f,** Microarray analysis of CDK8 expression in subtypes of 763 MB patient samples. n = 6 normal samples were collected by our institution. Mean ± SEM. Statistical analysis: Mann-Whitney U test. **g,** Microarray analysis of CDK8 expression in 4 subgroups of 763 MB patient samples. Mean ± SEM. Statistical analysis: Mann-Whitney U test. **h,** Immunoblot demonstrating the protein levels of c-MYC and CDK8 in D425 and D458 cells with genetic knockdown of CDK8. **i,** IVIS Spectrum signal of D458 xenografts with shCDK8 or shNull. Shown are days post injection.

**Extended Data Fig.2.**
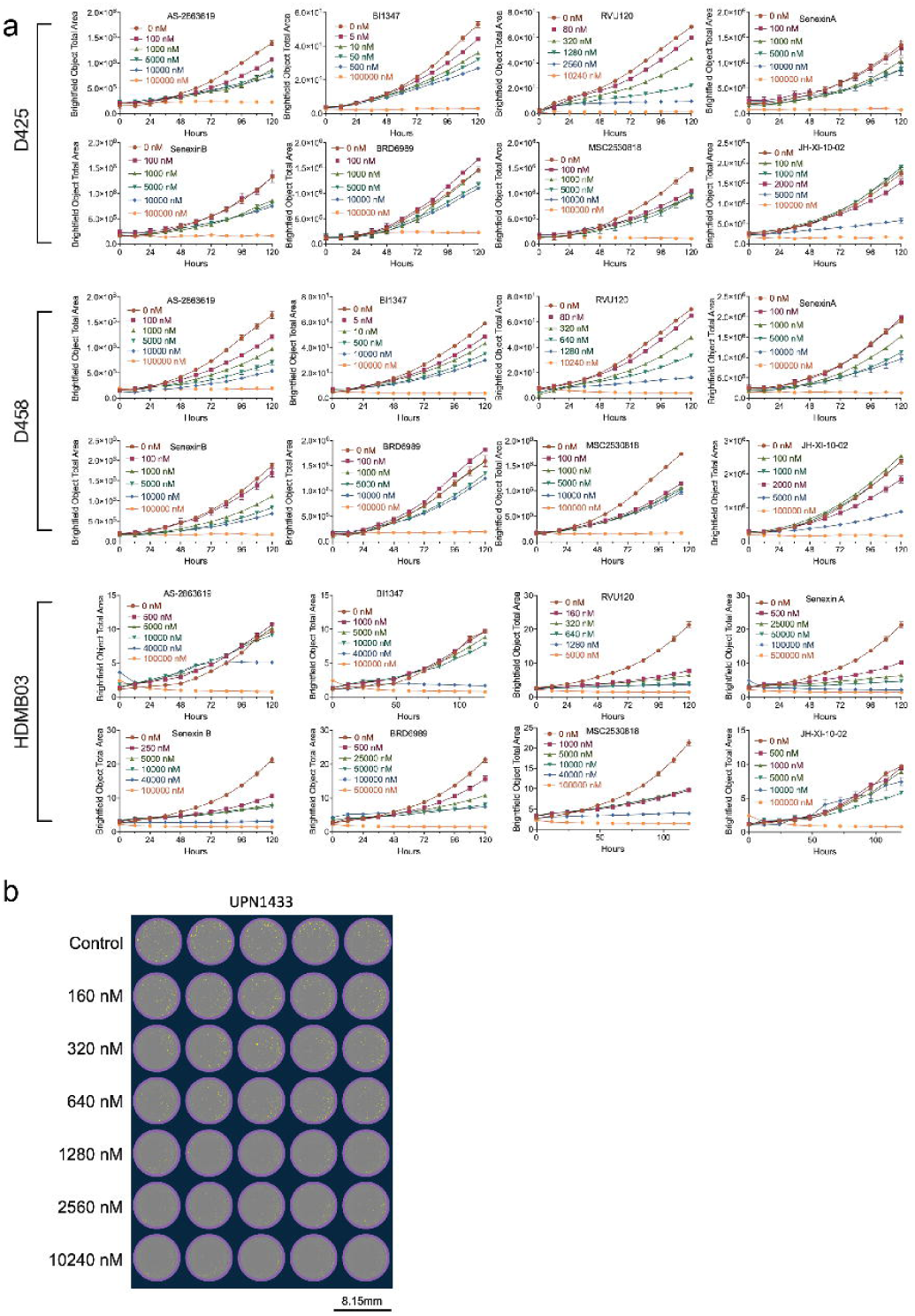
| The proliferation of medulloblastoma cells in the treatment of CDK8 inhibitors. **a**, Does response of CDK8 inhibitors treatment on D425, D458, and HDMB03 MB cells. The assays were performed in Incucyte Live-Cell Analysis System. **b,** Dose response of RVU120 treatment on human primary G3-MB cells.

**Extended Data Fig.3.**
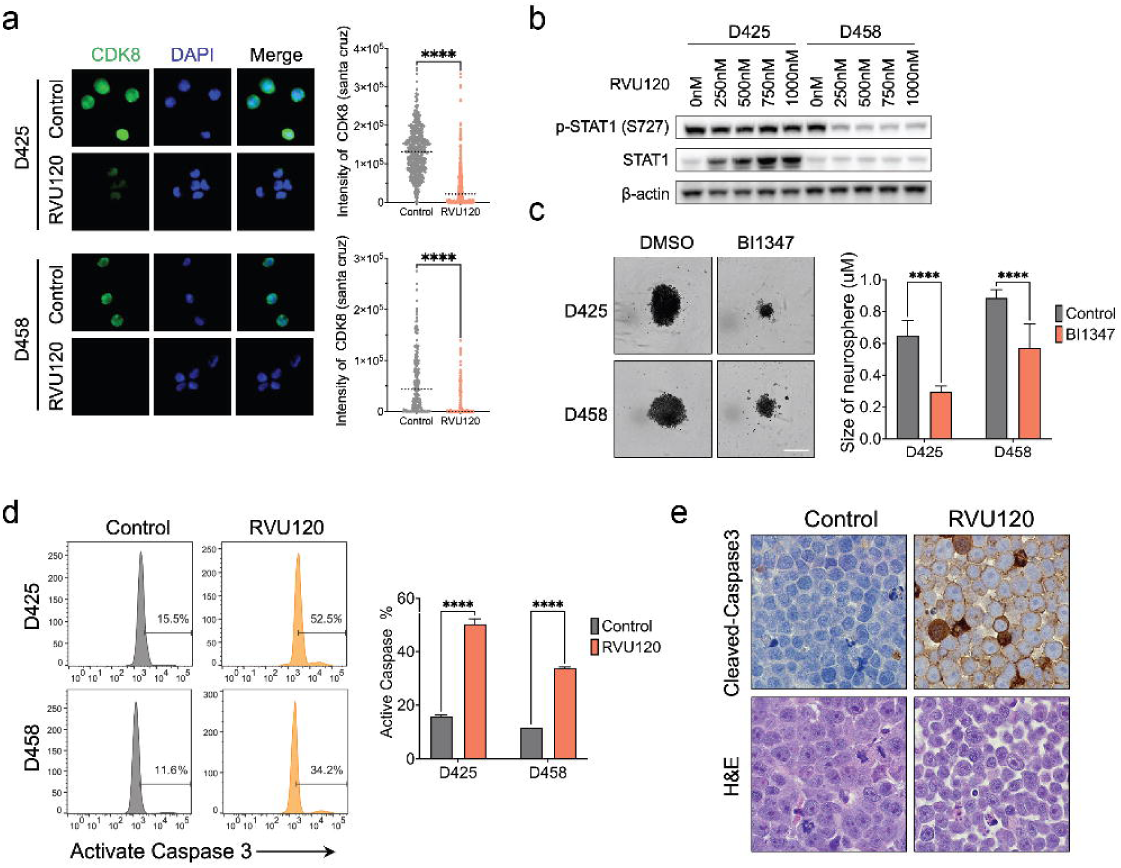
| RVU120 inhibits CDK8 activity in medulloblastoma cells. **a**, Immunofluorescence of CDK8 (green) and DAPI (blue) at 40X magnification. D425 and D458 cells were treated with IC50 RVU120 for 48 h. Mean ± SD. Scale bar, 10 μm. Statistical analysis: Mann-Whitney Wilcoxon test. **b,** Immunoblot demonstrating the protein levels of p-STAT1 (S727) and total STAT1 in D425 and D458 cells treated with various dose of RVU120. **c,** Representative images of neurosphere size in BI1347 treated D425 or D458 MB cell lines at 10 days are shown. n = 5. Mean ± SD. Scale bar, 400 μm. Statistical analysis: One-way ANOVA. **d,** Flow cytometric analysis of apoptotic population measured by active caspase 3. D425 and D458 cells were treated with IC50 RVU120 IC50 for 48 hours. Mean ± SD. Statistical analysis: One-way ANOVA. **e**, IHC analysis of cleaved-caspase3 in D458 xenograft mice treated with RVU120 compared to those treated with vehicle. Three mice in each group were treated for 10 days after tumor implantation. Original magnification, ×40.

**Extended Data Fig.4.**
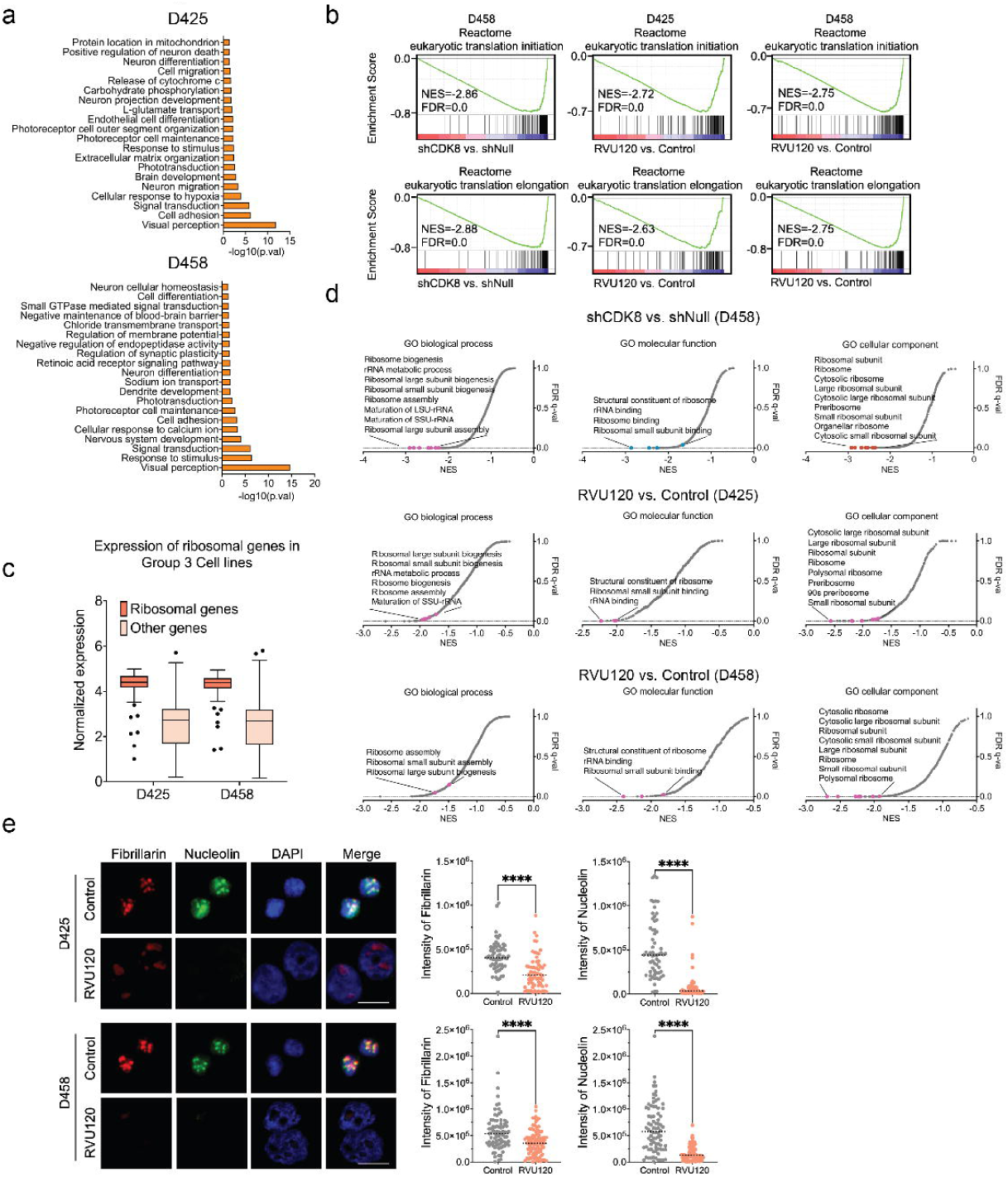
| CDK8 depletion leads to the repression of mRNA translation and ribosome biogenesis. **a**, Pathway enrichment analysis of D425 or D458 cells treated with RVU120. **b,** GSEA demonstrated downregulation of Reactome pathways related to mRNA translation initiation and elongation in genetic knockdown or pharmacological inhibition of CDK8. **c,** The normalized expression of ribosomal genes in D425 and D458 cells compared to all other genes. Mean ± SD. Statistical analysis: Kruskal-Wallis test. **d,** GSEA demonstrated downregulation of gene sets in three gene ontology categories related to ribosome biogenesis following genetic knockdown or pharmacological inhibition of CDK8. **e,** Immunofluorescence of Fibrillarin (red) , Nucleolin (green) and DAPI (blue) at 40X magnification. D425 and D458 cells were treated with IC50 RVU120 for 48 h. Mean ± SD. Scale bar, 10 μm. Statistical analysis: Mann-Whitney Wilcoxon test.

**Extended Data Fig.5.**
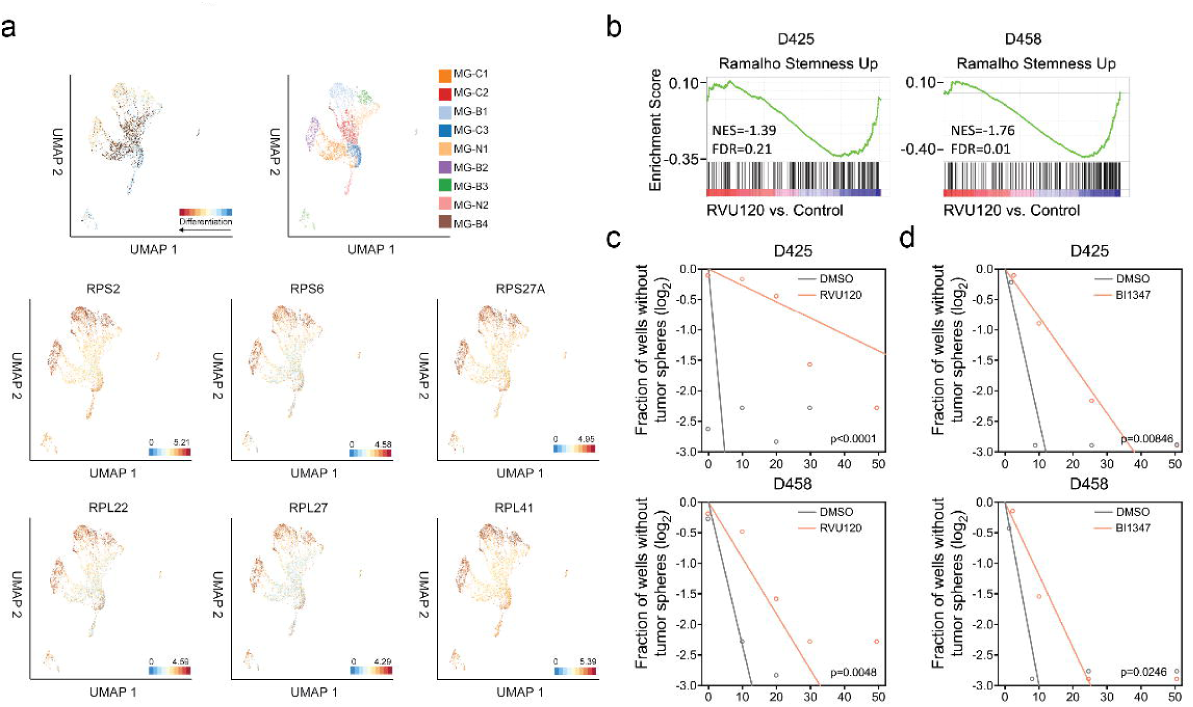
| CDK8 mediates stemness and differentiation in MYC-driven medulloblastoma. **a**, UMAP of 8144 cells of c-Myc and Gfi1 GP3 mouse models demonstrates differentiated cell and undifferentiated cell populations (top). Representative ribosomal gene expression is shown (bottom). **b,** GSEA demonstrated downregulation of the stemness pathway upon pharmacological inhibition of CDK8 in D425 and D458 MB cells. **c, d,** Sphere formation efficiency and self-renewal capacity were measured using extreme *in vitro* limiting dilution assays (ELDA) in two MB cell lines treated with either RVU120 or BI1347. P values were determined using the likelihood ratio test.

**Extended Data Fig.6.**
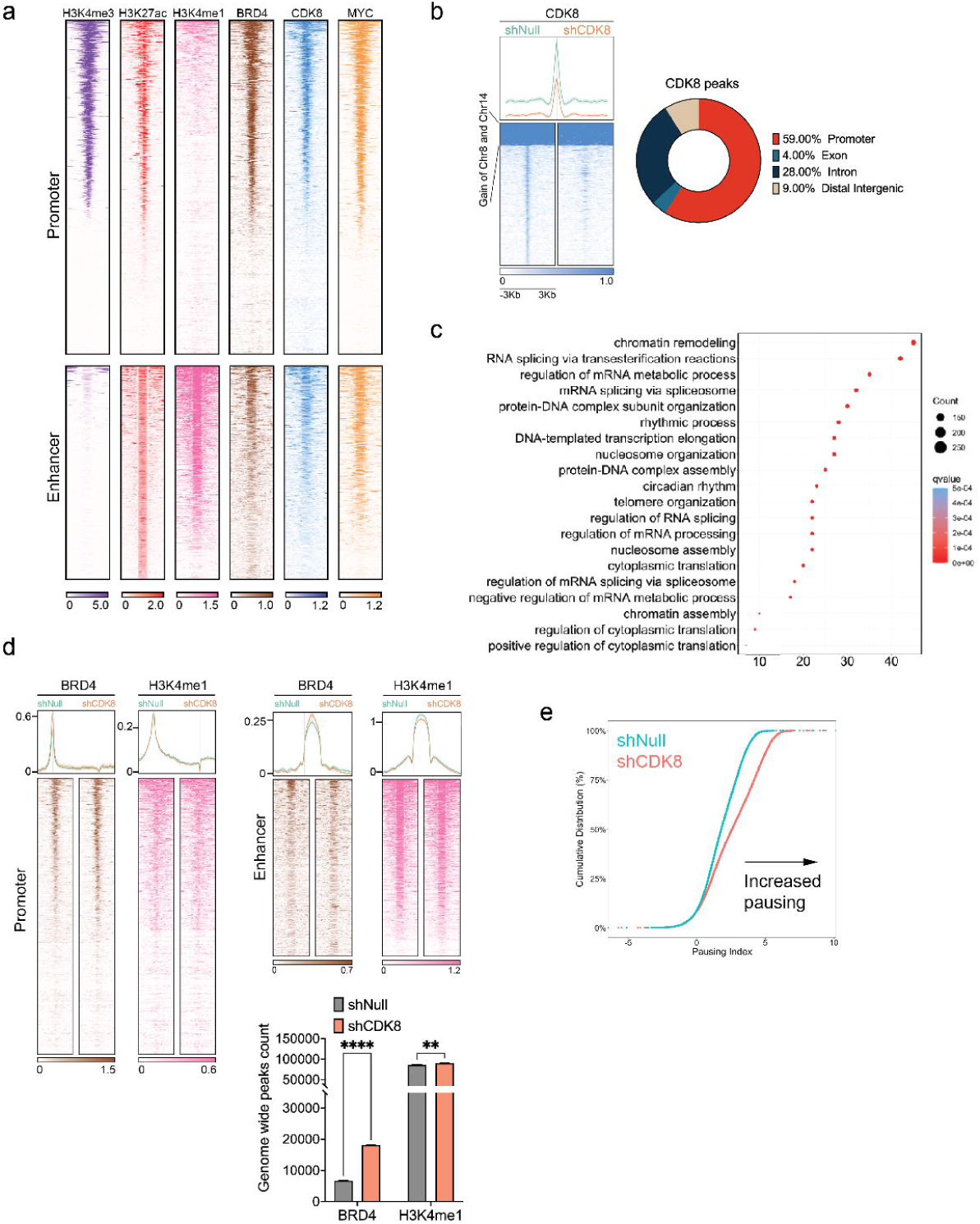
| CDK8 transcriptionally regulates ribosomal genes. **a**, Heatmaps showing CUT&RUN signals of CDK8, H3K4me3, H3K4me1, H3K27ac, BRD4, and MYC in D458 MB cell lines. The signals were displayed within a region spanning ± 3kb around the transcription start site (TSS). **b,** Heatmaps displaying genome-wide binding CUT&RUN signals of CDK8 in CDK8 knockdown D458 cells compared to control cells. The signals are displayed within a region spanning ± 3kb around the transcription start site (TSS). Pie chart showing the loss of CDK8 peaks were primarily localized at promoter sites. **c,** Pathway enrichment analysis identified mRNA translation related pathways as significantly enriched among genes exhibiting loss of CDK8 peaks following CDK8 knockdown. **d,** Bar plots and heatmaps showing CUT&RUN peak numbers and signals of BRD4 and H3K4me1 in D458 MB cells. The signals are displayed within a region spanning ± 3kb around the transcription start site (TSS). Mean ± SD. Statistical analysis: one-way ANOVA. **e,** Empirical cumulative distribution function (ECDF) plot shows significant increase in promoter-proximal pausing following CDK8 knockdown.

**Extended Data Fig.7.**
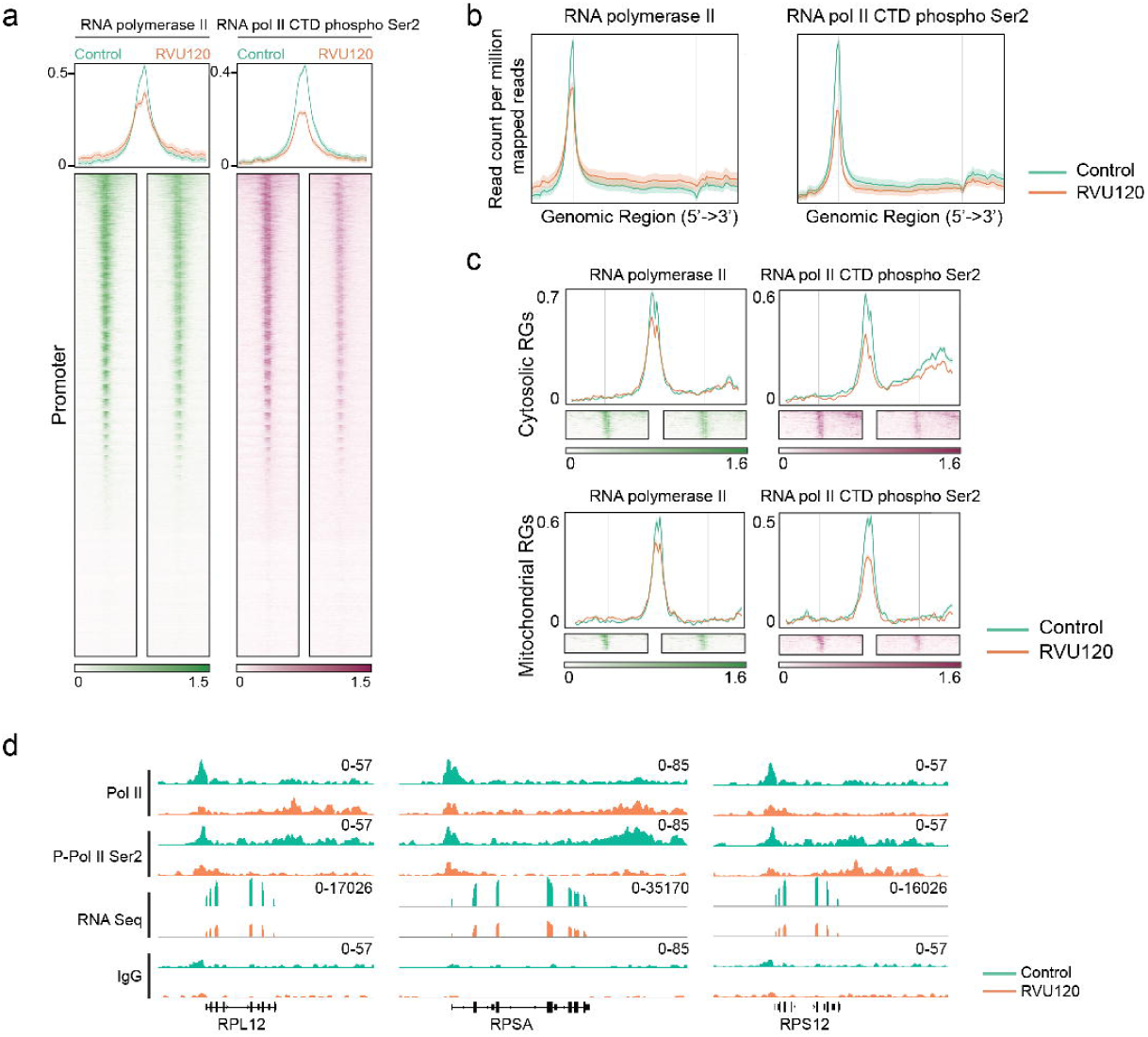
| Chromatin dynamic alterations in medulloblastoma cells treated with RVU120. **a**, Heatmaps showing CUT&RUN signals of RNA Pol II and phosphor-RNA Pol II in D458 MB cells treated with IC50 RVU120 for 48 hours. The signals are displayed within a region spanning ± 3kb around the transcription start site (TSS). **b,** Average distribution of RNA Pol II and phosphor-RNA Pol II peaks showing the alteration of RNA Pol II and phosphor-RNA Pol II signals across the gene body following the treatment of IC50 RVU120 for 48 hours. **c,** Average distribution and heatmaps of RNA Pol II and phosphor-RNA Pol II signals on cytosolic and mitochondrial ribosomal genes following the treatment of IC50 RVU120 for 48 hours. **d,** Representative examples of RNA Pol II and phosphor-RNA Pol II binding sites on ribosomal genes observed following the treatment of IC50 RVU120 for 48 hours.

**Extended Data Fig.8.**
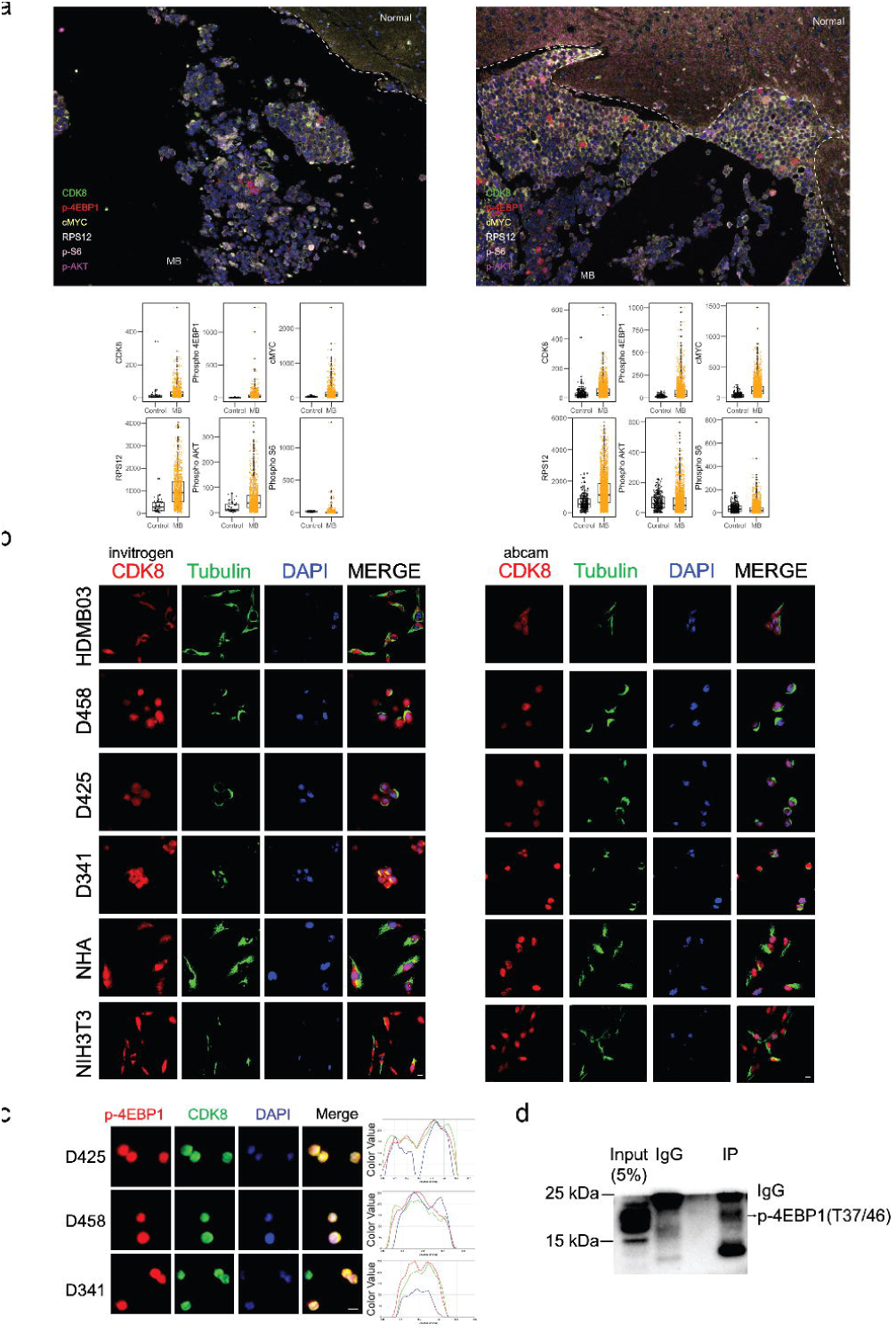
| Hyperactive ribosome biogenesis in MYC-driven Group 3 medulloblastoma. a, Multiplex IHC on G3-MB patient samples using CDK8, p-4EBP1, c-MYC, RPS12, p-S6, and p-AKT antibodies. p<0.05 in all biomarker groups. Statistical analysis: unparied t-test. **b,** Immunofluorescence analyses using two different CDK8 antibodies in MB cells, normal human astrocytes (NHA), and NIH3T3 cells. Scale bar: 10 μm**. c,** Immunofluorescence demonstrates the co-localization of p-4EBP1 (T37/46) and CDK8. Scale bar: 10 μm**. d,** The lysate from D458 cells was subjected to immunoprecipitation with CDK8 antibody, followed by the detection of co-precipitated p-4EBP1 using anti-p-4EBP1(T37/46) antibody.

**Extended Data Fig.9.**
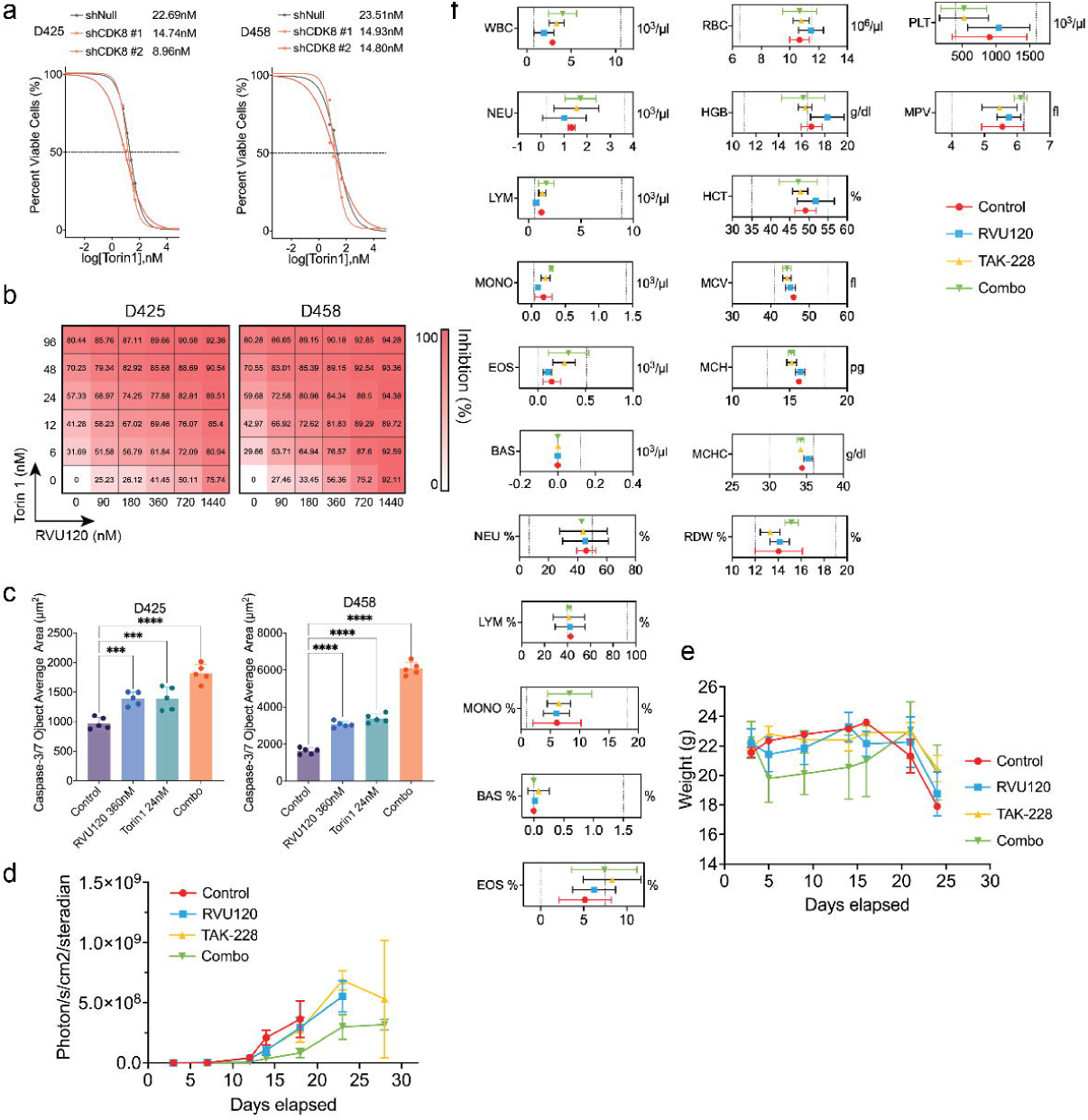
| Synergistic Targeting of CDK8 and mTOR in MYC-Driven medulloblastoma. **a**, IC50 determination of Troin1 in CDK8 knockdown D425 or D458 cells, with or without RVU120 treatment. **b,** Heatmap representation of inhibition percentage across a five-dose range of CDK8 inhibitor (RVU120) and mTOR inhibitor (Torin1) in D425 and D458 cells. Mean values of triple biological experiments are shown. **c,** The apoptosis assay was performed in Incucyte Nuclight Green-labeled MB cells treated with either DMSO, RVU120, Torin 1, or the combination of RVU120 and Torin1. Apoptotic cells were quantified using the caspase 3/7 Incucyte Live-Cell Analysis System. Mean ± SD. Statistical analysis: one-way ANOVA. **d,** IVIS Spectrum signal of D458 xenografts with treatment of either DMSO, RVU120, Torin 1, or the combination of RVU120 and Torin1. Shown are days post injection. **e,** Body weight of D458 xenografts with treatment of either DMSO, RVU120, Torin 1, or the combination of RVU120 and Torin1. Shown are days post injection. **f,** Hematological analysis of D458 xenograft mice. WBC: white blood cells, NEU: neutrophils, LYM: lymphocyte counts. MONO: monocytes. EOS: eosinophils. BAS: basophils. RBC: red blood cell count. HGB: Hemoglobin. HCT: Hematocrit. MCV: Mean Corpuscular Volume. MCH: Mean Corpuscular Hemoglobin. MCHC: Mean Corpuscular Hemoglobin Concentration. RDW: Red Cell Distribution Width. PLT: Platelet count. MPV: Mean Platelet Volume. Dotted lines represent the minimum or maximum of each parameter.

